# Regulation of ERK2 activity by dynamic *S*-acylation

**DOI:** 10.1101/2021.11.05.467491

**Authors:** Saara-Anne Azizi, Tian Qiu, Noah Brookes, Bryan C. Dickinson

## Abstract

The extracellular signal-regulated kinases (ERK1/2) are key effector proteins of the mitogen-activated protein kinase pathway, choreographing essential processes of cellular physiology. Critical in regulating these regulators are a patchwork of mechanisms, including post-translational modifications (PTMs) such as MEK-mediated phosphorylation. Here, we discover that ERK1/2 are subject to *S*-palmitoylation, a reversible lipid modification of cysteine residues, at C271/C254. Moreover, the levels of ERK1/2 *S*-acylation are modulated by epidermal growth factor (EGF) signaling, mirroring its phosphorylation dynamics, and palmitoylation-deficient ERK2 displays altered phosphorylation patterns at key sites. We find that chemical inhibition of either lipid addition or removal significantly alters ERK1/2’s EGF-triggered transcriptional program. We also identify a subset of “writer” protein acyl transferases (PATs) and an “eraser” acyl protein thioesterase (APT) that drive ERK1/2’s cycle of palmitoylation and depalmitoylation. Finally, we examine ERK1/2 *S*-acylation in a mouse model of metabolic syndrome, correlating changes in its lipidation levels with alterations in writer/eraser expression and solidifying the link between ERK1/2 activity, ERK1/2 lipidation, and organismal health. This study not only presents a previously undescribed mode of ERK1/2 regulation and a node to modulate MAPK pathway signaling in pathophysiological conditions, it also offers insight into the role of dynamic *S*-palmitoylation in cell signaling more generally.

## Introduction

Protein *S*-acylation, the addition of a long-chain fatty acid to a cysteine residue via a thioester bond, is a lipid post-translational modification (PTM) known to affect the activity and function of modified proteins. Protein *S*-acylation dynamics are enzymatically mediated, with DHHC domain-containing protein acyl transferases (PATs) acting as “writers” and acyl protein thioesterases (APTs) as “erasers” (Jiang et al., 2018). At the cellular level, these cycles of *S*-acylation and deacylation regulate protein subcellular trafficking and activity, and at the organismal level, have been implicated in cancer and neurological and inflammatory disease (Chamberlain & Shipston, 2015). One area in which the role of *S*-acylation has been increasingly recognized is signal transduction, with both receptors (MC1R, EGFR) and effectors (AKT, JNK, STAT3) requiring *S*-acylation for activity and downstream functionality (Blaustein et al., 2021; Chen et al., 2017; Runkle et al., 2016; Yang et al., 2012; Zhang et al., 2020). However, while the advent of chemical and biochemical techniques to study protein lipidation has precipitously increased our knowledge of the frequency and significance of *S*-acylation, the consequences of dynamic *S*-acylation for many proteins remains unknown.

The mitogen-activated kinases (MAPKs) are an evolutionarily conserved family of proteins that are critical in transducing extracellular signals to the interior of the cell and in regulating a diverse array of cellular programs (Cargnello & Roux, 2011; Morrison, 2012). Particularly well-studied within this venerable family are the extracellular signal-regulated kinases (ERK1/2). As the primary mediators of the Ras cascade, ERK1/2 phosphorylate hundreds of cytoplasmic and nuclear substrates, regulating processes such as embryogenesis, cell motility, proliferation, differentiation, and apoptosis (Chang et al., 2003; Lavoie et al., 2020; Sun et al., 2015). ERK1/2-dependent signal transduction is also critical in cellular and organismal metabolism, contributing to glycolytic pathway reprogramming (Warburg effect) in normal and cancer cells and to the development of metabolic syndrome (Papa et al., 2019).

The simple architecture of the central three-tiered cascade belies its complexity and finely tuned ability to determine specific cell responses. In fact, the master regulators ERK1/2 are themselves intricately regulated. Subcellular localization, docking motif-mediated interactions with substrates and regulatory elements, scaffolding proteins, phosphatases, and crosstalk with other signaling pathways – all modulate ERK specificity and activity (Ebisuya et al., 2005; Lake et al., 2016; Wortzel & Seger, 2011). Central to the ERK1/2 regulatory scheme are PTMs, in particular, phosphorylation. MEK1/2-mediated dual phosphorylation of a threonine-glutamic acid-tyrosine (TEY) motif is critical in regulating ERK1/2 activity following upstream signaling events. The magnitude and duration of phosphorylation at this motif determines the specific ERK1/2 transcriptional program. Other serine and threonine residues in ERK1/2 are also phosphorylated and regulate ERK1/2 activity (Berti & Seger, 2017; Chuderland et al., 2008; Oppermann et al., 2009). More recently, other PTMs have emerged as negative regulators of ERK1/2 activity, including the *S*-nitrosylation of Cys183, the acetylation of Lys72, and the tri-methylation of Lys302/361 of ERK1 (Feng et al., 2013; Vougiouklakis et al., 2015; Wu et al., 2018).

Despite its extensiveness, the established network of regulatory mechanisms does not fully explain how the activation of ERK1/2 by multiple extracellular stimuli results in distinct cellular outcomes, suggesting there is still more to be discovered about the regulation of ERK1/2 activity and function (Busca et al., 2016; Katz et al., 2007; Raman et al., 2007; Shaul & Seger, 2007). Intriguingly, proteomic studies suggest that ERK1/2 might be subject to *S*-acylation (Ren et al., 2013), leading us to hypothesize that lipidation may contribute to ERK1/2 regulation. Here, we show that ERK1/2 is *S*-acylated, in particular *S*-palmitoylated, *in cellulo* and *in vivo*. Moreover, we establish that ERK1/2 *S*-acylation levels are dynamically regulated during epidermal growth factor (EGF) stimulation and that *S*-acylation regulates the pattern of its TEY and serine phosphorylation and subsequent activation. We then show ERK1/2 *S*-acylation is enzymatically mediated by dynamic associations with a subset of DHHC-PATs, which install the acyl group, and APT2, which removes the acyl group. Finally, we use a mouse model of metabolic syndrome to profile changes in *S*-acylation correlated to aberrant activity and changes in writer and eraser protein expression. This work introduces *S*-acylation as a novel regulatory module of ERK1/2 activity and potential therapeutic node for targeting ERK1/2 activity, not only in metabolic syndrome but in cancer and other MAPK signaling-driven pathologies.

## Results

### ERK1/2 *S*-acylation levels respond to EGF stimulation

ERK1/2 acylation has been hinted by thiopropyl capture carried out in adipocytes (Ren et al., 2013). Therefore, we sought to confirm that *S*-acylation, and more specifically, *S*-palmitoylation, is a PTM of ERK1/2. To visualize cysteine acylation, we first used acyl biotin exchange (ABE), in which the labile thioester bond of acylated cysteine is cleaved by hydroxylamine (HA) and substituted with an enrichment handle (Drisdel & Green, 2004). This assay, which was carried out in HEK293T, A431, and HepG2 cells, revealed that ERK1/2 are *S*-acylated across cell lines (Figure 1A). Next, to further corroborate these results and to confirm the incorporation of palmitate, we used metabolic labeling with 17-octadecynoic acid (17-ODYA), the ω-alkyne analogue of the 16C palmitic acid. Here, we observed incorporation of the tagged palmitic acid in HEK293T cells, affirming ERK1/2 *S*-palmitoylation (Figure 1B) (Charron et al., 2009). Importantly, hydroxylamine (HA) treatment diminished the signal in this experiment, confirming thioester bond formation and thus cysteine modification (Figure 1B). We also probed ERK *S*-acylation by ABE in mouse tissues, observing significant *S*-acylation across all tissues profiled, especially in the brain, liver, and pancreas, confirming ERK *S*-acylation *in vivo* (Figure 1C). Together, these results establish that ERK1/2 are *S*-acylated at the cellular and organismal levels.

**Figure 1.**
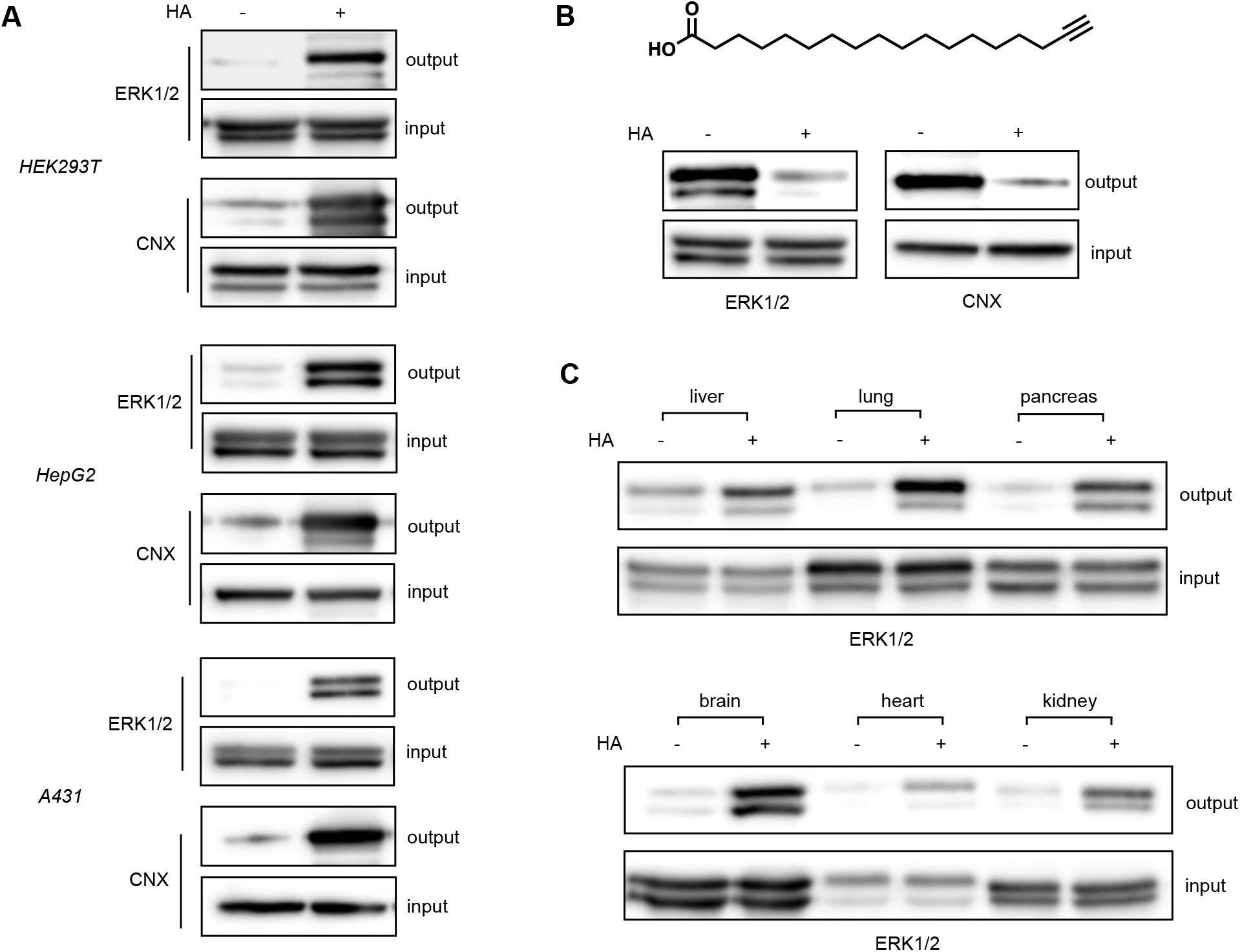
ERK1/2 are *S*-palmitoylated. (**A**) Acyl biotin exchange (ABE) assay carried out in HEK293T, A431, and HepG2 cells, wherein signal in the hydroxylamine (HA) treated samples indicates *S*-acylation. Calnexin (CNX) is used as loading and assay control. (**B**) Metabolic labeling with 17-octadecanoic (17-ODYA) in HEK293T cells. Signal in ‘-HA’ lanes indicates the incorporation of the 16C fatty acid, while loss of signal in the ‘+HA’ samples indicates cysteine bond formation. (**C**) ABE carried on a panel of tissues harvested from C57BL/6 mice to visualize *S*-acylation *in vivo*.

Central to ERK1/2’s mediation of cellular events is their ligand-induced activation, which stimulates a rapid increase in their phosphorylation and then downstream kinase activity. We next sought to determine whether the levels of ERK1/2 *S*-acylation are sensitive to ligand-induced activation, as changes in ERK1/2 *S*-palmitoylation concurrent with activity changes could hint at a regulatory role for *S*-palmitoylation in ERK1/2 activation. We therefore monitored ERK1/2 *S*-acylation levels upon stimulation with the epidermal growth factor (EGF), a secreted peptide which activates a signal transduction network that includes the RAS/ERK pathway (Katz et al., 2007). Upon treatment of serum-starved cells with EGF, the basal *S*-acylation of ERK1/2 increases within 5 minutes and remains elevated through 15 minutes of stimulation, with a return to baseline acylation levels occurring by 30 minutes in both HepG2 and A431 cells (Figure 2A, S1A,B). Significantly, this pattern mirrors that of ERK phosphorylation, which also remains elevated through 15 minutes and decreases by 30 minutes (Kiyatkin et al., 2020). Since much remains unknown about how the extracellular concentration of ligand (signal strength) is relayed intracellularly, we next assessed the dose-dependence of ERK1/2’s *S*-acylation increase. As EGF concentration increased from 0 to 1 to 10 ng mL^−1^, ERK1/2 *S*-acylation increased in-step, with a lessening increase at 100 ng mL^−1^ in HepG2 and A431 cells. This suggests that *S*-acylation is responsive to the flow of quantitative information through the pathway (Figure 2B, S1C,D). These results established our activating concentration of EGF at 1 ng mL^−1^, a concentration congruent with reported physiologic levels of the growth factor in some tissues (Pinilla-Macua et al., 2017). Finally, to assess the generality of the ERK *S*-acylation dynamics, we probed for ERK1/2 *S*-acylation changes upon treatment with insulin, another signaling ligand also established to activate the ERK/MAPK pathway. To our surprise, insulin treatment did not trigger an increase in ERK1/2 *S*-acylation in either HepG2 or A431 cells, although a decrease in *S*-acylation was observed in A431 cells at 30 minutes (Figure 2C, S1E). Thus, the *S*-acylation of ERK1/2 is not only dynamic, but also sensitive to signal strength and signal identity, emphasizing its potential as a regulator of ERK1/2 activity.

**Figure 2.**
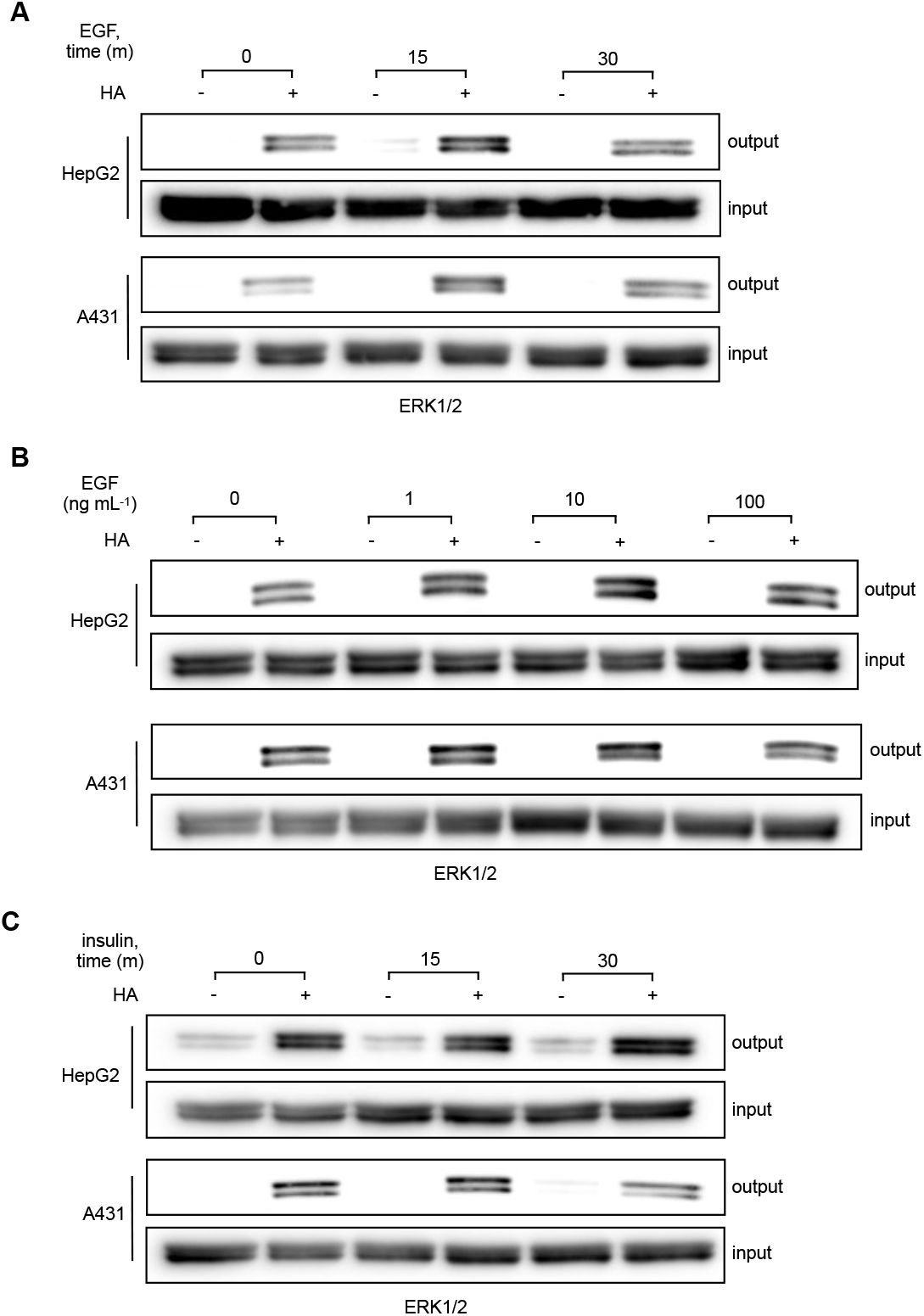
ERK1/2 *S*-acylation responds to EGF stimulation. (**A**) ABE assay carried out in HepG2 and A431 cells following stimulation with EGF (1 ng mL^−1^) for 0, 15, and 30 minutes. (**B**) ABE assay carried out in HepG2 and A431 cells following stimulation with 0, 1, 10, and 100 ng mL^−1^ EGF for 15 minutes. (**C**) ABE assay carried out in HepG2 and A431 cells following stimulation with insulin (10 nM) for 0, 15, and 30 minutes.

### Dynamic ERK *S*-acylation does not depend on TEY motif phosphorylation

Given that the timing of ERK1/2’s dynamic *S*-acylation corresponds to that of its dynamic TEY phosphorylation, we next attempted to parse the connection between the two modifications. First, we used ABE to determine the *S*-acylation status of active ERK1/2, i.e., ERK1/2 that have been phosphorylated at the TEY motif of their activation loops. Here, we observed palmitoylated ERK1/2 increasing concomitantly with ERK1/2 phosphorylation (Figure 3A). This result confirms that the two PTMs can co-occur on ERK1/2. We next used established MAPK pathway inhibitors to investigate whether the signaling elements that regulate phosphorylation also regulate ERK1/2’s dynamic palmitoylation. Pre-treatment with upstream inhibitors for MEK (AZD6244, CI-1040) and RAF (PLX-4720), as well as with okadaic acid, an inhibitor of ERK1/2-directed phosphatase activity, inhibited the EGF-induced increase in ERK1/2 *S*-acylation (Figure S2A,B) (Cohen et al., 1989; Sebolt-Leopold et al., 1999; Tai et al., 2016; Tsai et al., 2008). These results suggest that intact regulatory infrastructure is necessary for the dynamic *S*-acylation of ERK1/2. However, while these data establish that the dynamic *S*-acylation of ERK1/2 is connected to pathway activation, they do not reveal whether ERK1/2 phosphorylation itself is necessary for *S*-acylation.

**Figure 3.**
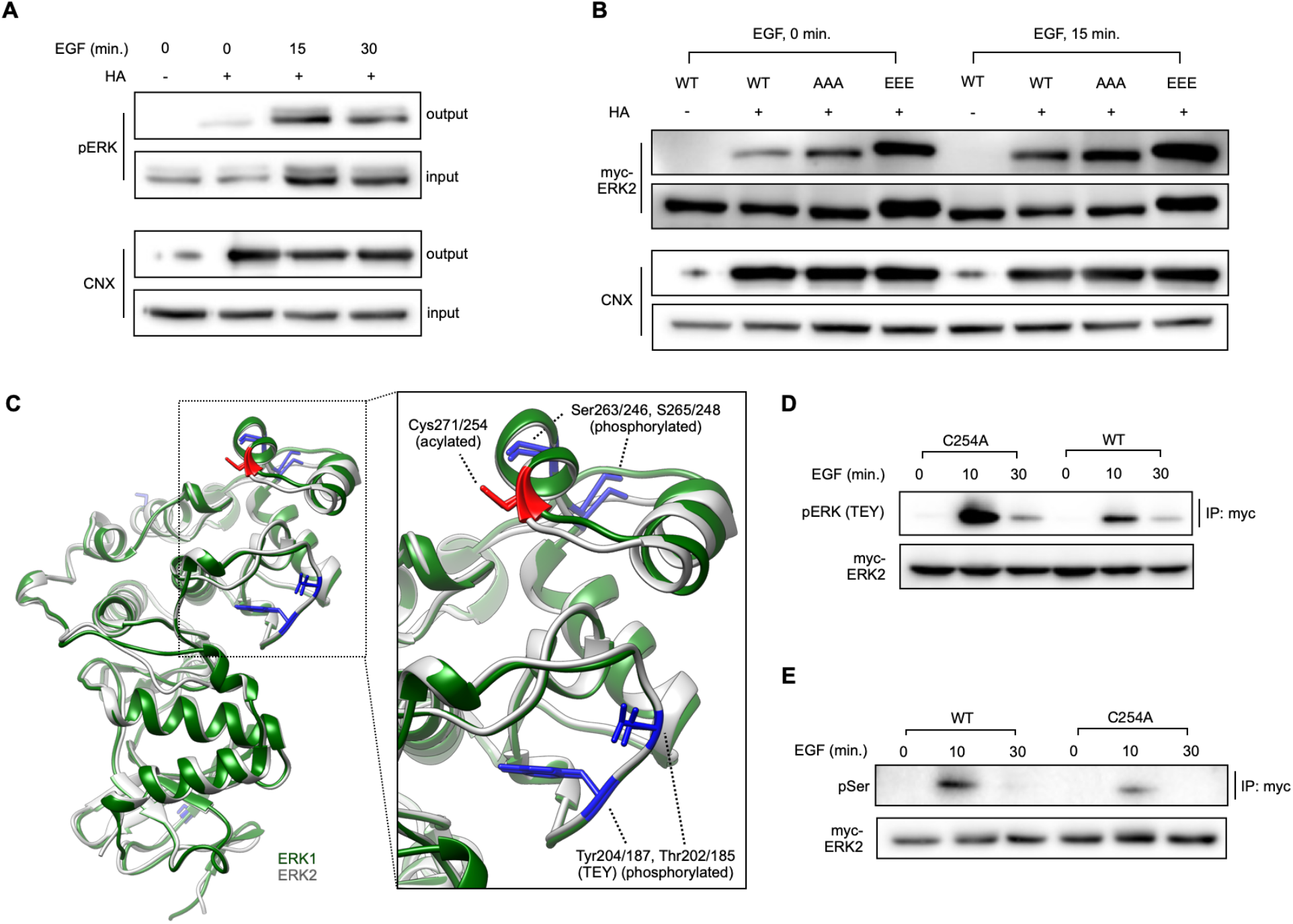
Mapping crosstalk between ERK2 *S*-acylation and phosphorylation. (**A**) ABE assay carried out in HepG2 cells and analyzed for phospho ERK1/2 (Thr185/202,Tyr187/204) via Western blotting. (**B**) ABE assay carried out in HEK293T cells overexpressing WT, TEY(AAA), or TEY(EEE) ERK2 and stimulated with EGF (1 ng mL^−1^) for 15 minutes. (**C**) Crystal structure of the C-terminal lobes of ERK1 and 2, demonstrating structural homology, as well as phosphorylation (blue) and acylation (red) sites. (**D**) Immunoprecipitation of myc-ERK2 (WT or C254A) in HEK293T cells, with or without EGF stimulation (10 and 30 minutes, 1 ng mL^−1^) and analyzed via Western blotting for phospho-ERK (Thr185/Tyr187). (**E**) Immunoprecipitation of myc-ERK2 (WT, C254A), with or without EGF stimulation (10, 30 minutes, 1 ng mL^−1^), analyzed via Western blotting for phosphoserine.

In order to gain further insight into the sequentiality and interdependence of the two modifications, we created variants of ERK2 – the more widely and abundantly expressed isoform of ERK – in which the three sites of the phospho-acceptor motif were mutated to either alanine (AAA) or glutamic acid (EEE). These mutations respectively confer resistance to or mimic constitutive phosphorylation, enabling us to probe the result of perturbing ERK2 phosphorylation on its palmitoylation. Overexpression of these constructs in HEK293T cells, followed by EGF stimulation and visualization of acylated proteins, indicated that ERK2’s dynamic *S*-acylation was not disrupted in either the constitutively inactive or active constructs (Figure 3B). This suggests that the EGF-mediated increase in ERK2’s *S*-acylation is not dependent on its phosphorylation status. Interestingly, the levels of basal *S*-acylation in serum-starved cells increased for both constructs relative to the wild type (WT), again hinting at a more complicated regulatory network (Figure 3B). Collectively, these results demonstrate that dynamic ERK1/2 *S*-palmitoylation is dependent on pathway activation but remains intact when the phosphorylation of ERK is disrupted.

### Palmitoylation-deficient ERK2 displays altered phosphorylation patterns

To determine the consequences of *S*-acylation for the regulation and activation of ERK1/2, we next sought to generate palmitoylation-deficient variants of each ERK. This required mapping the *S*-acylation site(s), wherein we individually mutated each of the six cysteines of ERK1 and the seven cysteines of ERK2 to serine or alanine, and then used ABE to visualize the *S*-acylation status of each C-to-S mutant. For ERK1, we observed a decrease in ABE signal for both C239S and C271S, and for ERK2, a decrease was seen for both C216S and C254S/C254A – signifying that these cysteines are sites of *S*-acylation (Figure S3A-C). As the loss of signal for C233 in ERK1 and C216 in ERK2 were accompanied by a significant loss of expression for these mutants, implicating these residues (and possibly its *S*-acylation) in the stability or proteasome degradation of ERK1/2, we chose to focus on the *S*-acylation of C254 and C271. Importantly, these residues are located in the C-terminal lobe of ERK1/2, which contains their activation and catalytic loops and is also subject to other regulatory PTMs (Peti & Page, 2013).

With these biochemical tools in hand, we next assessed the impact of ERK1/2 *S*-acylation on their activation, as measured by TEY phosphorylation. For this, we focused on ERK2, given its expression exceeds that of ERK1 in most cells and that the two kinases possess highly similar sequences and identical substrate specificity *in vitro*. Overexpression of myc-tagged WT and C254A ERK2, followed by EGF stimulation and immunoprecipitation of the epitope tagged-constructs, allowed for visualization and comparison of TEY phosphorylation of the WT vs. *S*-acylation-deficient ERK, while eliminating background signal from endogenous ERK1/2. Strikingly, myc-ERK2(C254A) was dramatically more phosphorylated at the TEY motif than its WT counterpart (Figure 3C, S3D). This TEY hyperphosphorylation upon EGF stimulation lost upon inhibition of MEK1/2 (Figure S3E) (Kutzleb et al., 1998). Next, as the site of *S*-acylation, C254, is most proximal not to the Thr185/Tyr187 phosphorylation sites, but rather to the purportedly phosphorylated serine residues Ser246/248, we assayed for serine phosphorylation under the same conditions (Figure 3B). Here, we found that in contrast to the TEY phosphorylation, myc-ERK2(C254A) serine phosphorylation was diminished relative to that of WT ERK2 (Figure 3D). Together, these results suggest that the *S*-acylation of C254 has a substantial role in regulating the phosphorylation patterns, and we hypothesize, the activation of ERK2.

### Chemical inhibition of ERK1/2 *S*-acylation disrupts its transcriptional program

Upon EGF stimulation and translocation to the nucleus, ERK signaling elicits a gene expression program linked to diverse cellular events, such as proliferation and differentiation. Given the significant but opposing changes in TEY and SPS phosphorylation, we next aimed to determine how loss of dynamic ERK1/2 *S*-acylation affects this transcriptional program. To do this, we began by identifying chemical tools to disrupt the cycle of ERK1/2 *S*-acylation and deacylation, as overexpressed proteins have the potential to overload biological pathways (Bolognesi & Lehner, 2018). First, we targeted the installation of the lipid by the DHHC-PATs using both 2-bromopalmitate (2-BP), a widely used albeit promiscuous inhibitor, and cyano-myracrylamide (CMA), another small inhibitor with a different reactivity profile recently developed by our group (Azizi et al., 2021; Lan et al., 2021; Webb et al., 2000). ABE revealed that treatment with 2-BP (20 μM, 6 hours) failed to decrease the *S*-acylation of ERK, possibly due to its poor potency and/or poor targeting of ERK-specific writers (Figure 4A). However, treatment with CMA, but not its inactive analogue, under parallel conditions (20 uM, 6 hours) resulted in a significant decrease of the *S*-acylation of ERK1/2, with maximum inhibition of *S*-acylation reached after three hours of treatment in HepG2 cells (Figure 4A, S4A,B). These results validate CMA as a tool to analyze the downstream consequences of inhibiting ERK1/2 *S*-acylation and confirm that the *S*-acylation of ERK1/2 are enzymatically regulated.

**Figure 4.**
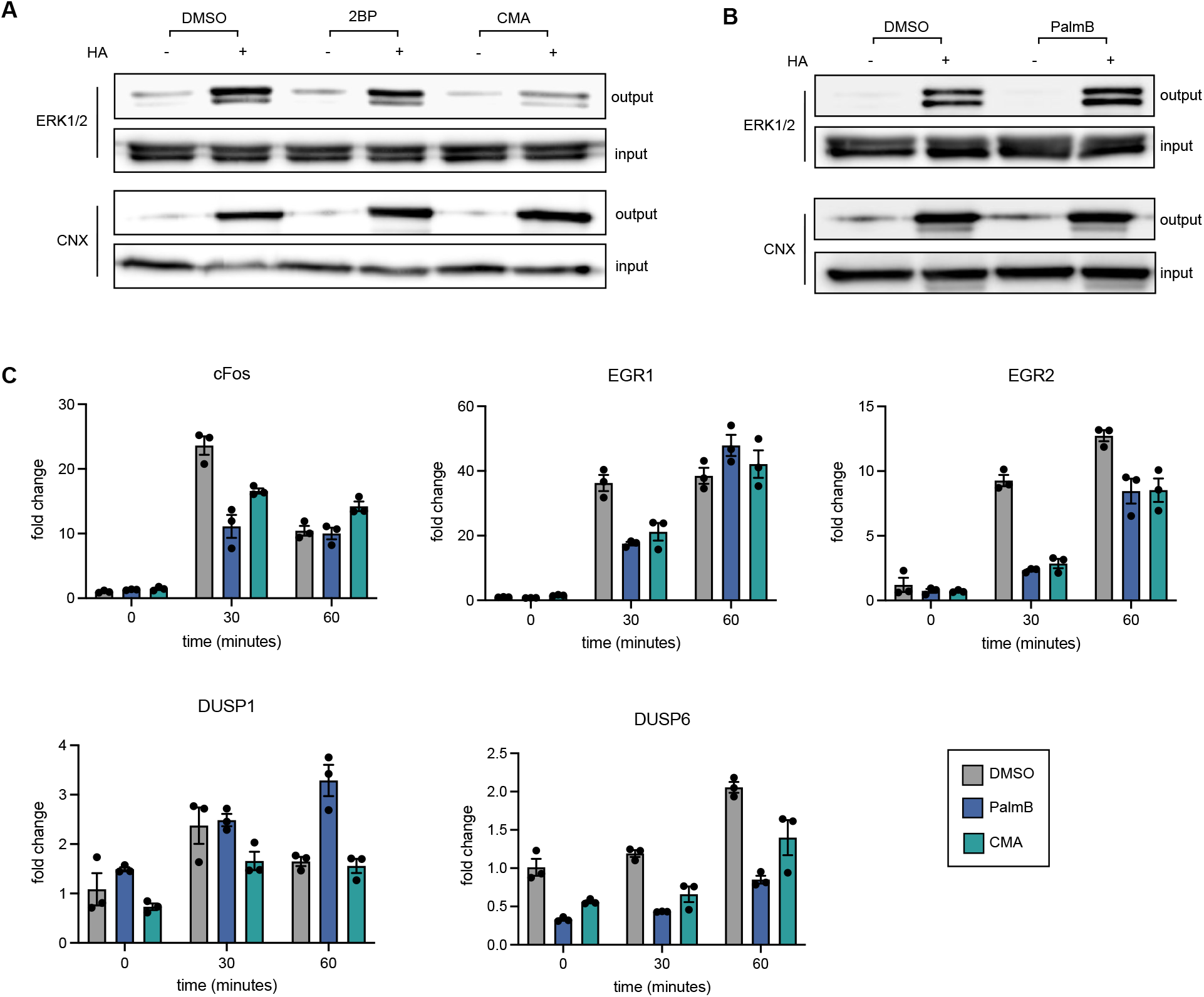
Chemical perturbation of ERK1/2 *S*-acylation disrupts their transcriptional program. (**A**) ABE assay carried out in HepG2 cells treated with DMSO, 2BP (20 μM), or CMA (20 μM) for 6 hours. Calnexin (CNX) is used as loading and assay control. (**B**) ABE assay carried out in HepG2 cells treated with DMSO or PalmB (20 μM) for 6 hours. Calnexin (CNX) is used as loading and assay control. (**C**) RT-qPCR of key transcripts in the EGF-stimulated transcriptional program in HepG2 cells treated with DMSO, CMA (20 μM), or PalmB (20 μM) for three hours, followed by stimulation with EGF (1 ng mL^−1^) for 30 and 60 minutes.

Next, we used palmostatin B (PalmB), a pan inhibitor of the APTs, to interrupt enzymatic lipid removal, and therefore increase the *S*-acylation of ERK1/2 (Dekker et al., 2010). Treatment of HepG2 cells with PalmB (20 μM, 6 hours) and visualization of acylated proteins via ABE showed increased ERK1/2 *S*-acylation, confirming the utility of this inhibitor in probing the effects of perturbing ERK1/2 *S*-acylation (Figure 4B). The effectiveness of PalmB also confirmed that ERK1/2 deacylation is enzymatically mediated. We next sought to determine which APT(s) are responsible for the deacylation of ERK1/2. While the β-lactone PalmB acts on globally on APTs, ML-348 and ML-349 are inhibitors specific for two of the most widely studied *S*-deacylases, APT1 and APT2, respectively (Adibekian et al., 2010a, 2010b). We found that an increase in ERK *S*-acylation only occurred with ML-349 treatment (Figure S4C), suggesting that APT2, but not APT1, regulates the deacylation of ERK1/2. However, as the increase was less than that observed with PalmB, it also introduces the possibility that additional APTs have a role in the regulation of ERK1/2 and establishes PalmB as a key tool to disrupt ERK1/2’s deacylation.

With these validated chemical inhibitors of ERK1/2 *S*-acylation and deacylation in hand, we next aimed to assess the consequences of disrupted *S*-acylation for ERK1/2 transcriptional regulation. As the key activator of MAPK pathway, ERK1/2 is connected to the induction of hundreds of gene targets, including cFos, early growth response proteins 1/2 (EGR1/2), and the dual specificity phosphatases 1/6 (DUSP1/6) (Uhlitz et al., 2017). Reverse transcription-quantitative PCR (RT-qPCR), following pretreatment of HepG2 cells with DMSO, CMA, or PalmB and stimulation with EGF (0, 30, and 60 minutes), revealed inhibitor-dependent changes in the induction pattern of these genes (Figure 4C). In the case of the immediate early response genes cFos and EGR1/2, treatment with both CMA and PalmB limited the rapid mRNA transcription typically observed after 30 minutes of EGF treatment, as compared to the DMSO control. However, after 60 minutes, cFos and EGR1 transcript levels matched the control sample, while EGR2 transcript levels continued to rise. This suggests that disruption of ERK1/2 *S*-acylation – both its installation and removal – delays rather than inhibits its EGF-stimulated transcriptional program. For the immediate late gene DUSP1, PalmB treatment resulted in increased activation at 60 minutes, while DUSP6 expression was dampened at all time points. Together, these alterations suggest that the cycle of *S*-acylation is significant in regulating both the timing and amplitude of ERK-mediated gene expression changes.

### ERK2 *S*-acylation is mediated by dynamic associations with DHHCs and APTs

Having determined that dynamic *S*-acylation is significant for ERK1/2 activity, we next sought to identify the molecular determinants of this PTM. Identification of these regulatory elements, i.e., the writers and erasers, could enable more controlled modulation ERK *S*-acylation and therefore represents a novel way to manipulate ERK1/2 activity. Typically, the “Fukata assay” – overexpression of 23 mammalian zDHHCs (Fukata et al., 2006), followed by visualization of *S*-acylation via ABE – is used to ascertain the DHHC-PAT writers of a particular substrate. However, in this screen, while slight increases in ERK1/2 *S*-acylation were observed with overexpression of some zDHHCs, no definitive writers for ERK1/2 were identified (Figure S5A). This could suggest that ERK1/2 *S*-acylation is regulated by multiple PATs, with compensatory down-regulation complicating observation of increases in *S*-acylation. In addition, the levels of ERK1/2 *S*-acylation are likely tightly regulated, and the ABE assay reveals the steady-state levels of *S*-acylation for a target, not the cycling of *S*-acylation mediated in part by the PATs. Therefore, we next used metabolic labeling to visualize a change in the rate of *S*-acylation via increased incorporation of an alkyne tagged fatty acid. Here, we observed that a panel of PATs – annotated as zDHHC2, 3, 7, 9, 11, 23, 12, 14, 17, 20, and 21 – all increased ERK1/2 palmitoylation, with zDHHCs 7, 11, 14, 17, 20, 21 being the strongest hits for both ERK1 and ERK2 (Figure 5A).

**Figure 5.**
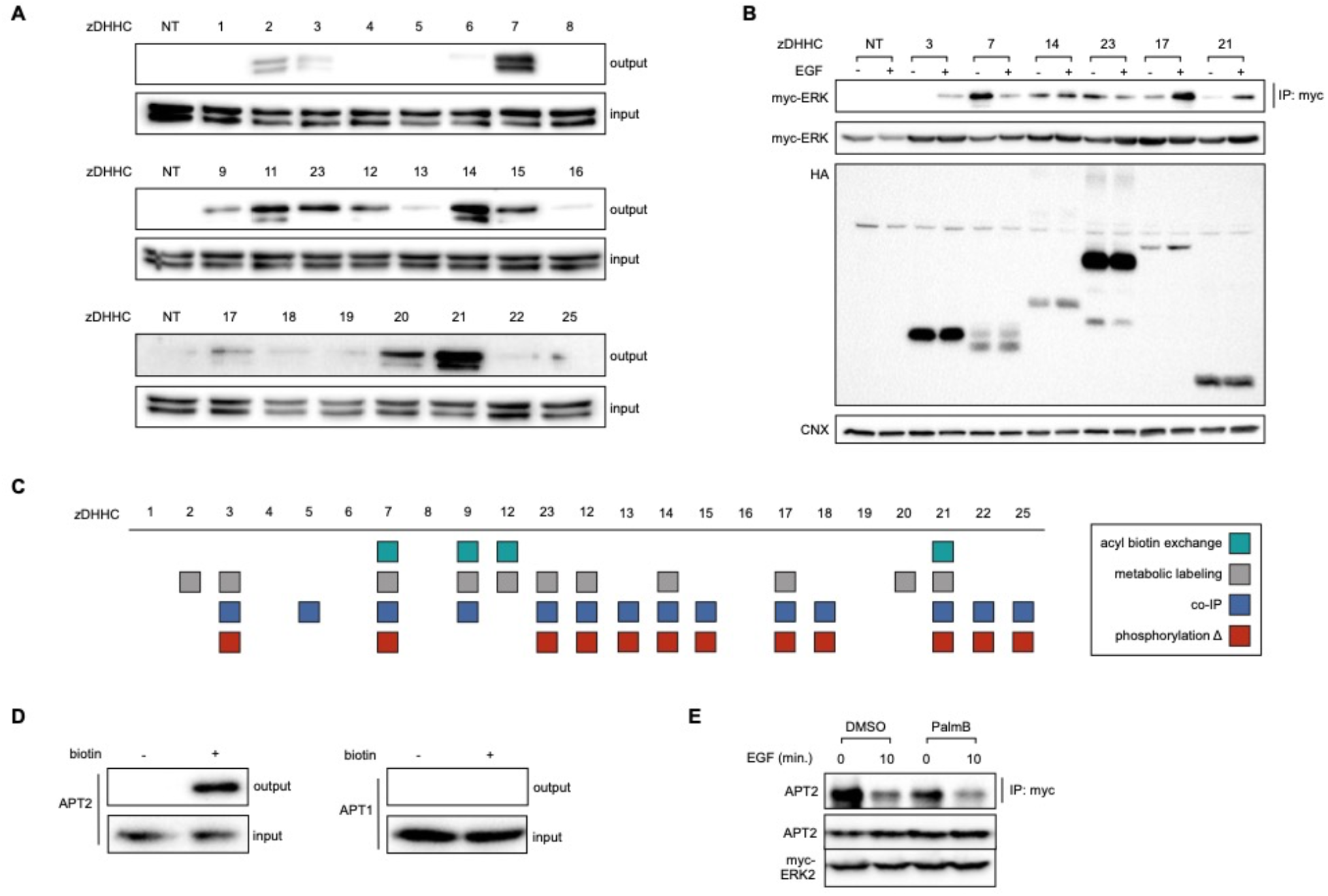
ERK2’s acylation is mediated by dynamic associations with writer and eraser proteins. (**A**) ERK1/2 metabolic labeling with 17-ODYA in HEK293T cells overexpressing murine DHHC family proteins. (**B**) Co-immunoprecipitation of myc-ERK2 with indicated HA-tagged DHHC proteins in HEK293T cells with or without EGF stimulation (10 minutes, 1 ng mL^−1^). Co-immunoprecipitated proteins were visualized via Western blotting for myc-ERK2. (**C**) Diagram summarizing the DHHC “hits” from the four assays (ABE, metabolic labeling, TEY phosphorylation changes, and co-immunoprecipitation) used to ascertain ERK2 writers. (**D**) Overexpression of myc-ERK2 tagged with TurboID in HEK293T cells, followed by biotin incubation and streptavidin enrichment of labeled proteins. Enriched proteins were visualized via Western blotting for APT1 and APT2. (**E**) Co-immunoprecipitation of APT2 with myc-ERK2 in HEK293T cells with or without EGF stimulation (10 minutes, 1 ng mL^−1^) and with or without PalmB treatment (20 μM, 3 hours). Co-immunoprecipitated proteins were visualized via Western blotting for APT2.

Given that metabolic labeling implicated a large number (11/23) DHHC proteins as ERK1/2 writers, we next aimed to determine which were significant in its EGF-promoted dynamic *S*-acylation. Having observed a change in the TEY phosphorylation of palmitoylation deficient ERK2, we next probed for changes in the ERK2 EGF-stimulated TEY phosphorylation concurrent with DHHC overexpression. In this screen, we observed that a number of DHHCs (3, 7, 23, 12, 13, 14, 15, 17, 18, 21, 22, and 25) increased ERK2 TEY phosphorylation, with strong overlap (7/12) with writers identified via metabolic labeling (Figure 5C, S5B). For final validation, we also assessed the physical writer/substrate interaction in the context of EGF signaling, via co-immunoprecipitation of ERK2 by HA-tagged zDHHC family proteins. Here, a panel of DHHCs (3, 5, 7, 9, 11, 14, 17, 18, 20, 21, 22, and 23) interacted with ERK2 (Figure 5B-C, S5C). Summation of the results confirmed DHHC7 and 21 – hits in all assays – as key writers of ERK2 *S*-acylation. Significantly, ERK2 association with certain PATs was dynamic, with ERK2 dissociating from zDHHCs 5, 7, and 23 and associating with zDHHCs 3, 17, and 21 upon stimulation with EGF (Figure 5B, S5C). These results demonstrate that certain DHHC family members are not just writers of ERK2 *S*-acylation, they are also central to its dynamic *S*-acylation during EGF signaling.

The other vital element in the cycle of *S*-acylation/deacylation are the erasers, the APTs (Azizi et al., 2019; Cao et al., 2019; Lin & Conibear, 2015). To identify candidate erasers, we first overexpressed the cytosolic serine hydrolases annotated as APTs, APT1, APT2, and ABHD17A/B/C, and assayed for resultant changes in ERK1/2 *S*-acylation. Here, no discernible decreases – which would indicate ERK1/2-targeted eraser activity – were observed (Figure S5D). Since previous experiments with the inhibitor ML-349 intimated that APT2 was a candidate eraser, we next decided to see if knockdown of this protein (encoded by LYPLA2) would elicit a change in ERK1/2 S-acylation. However, neither siRNA mediated knockdown of APT2 or APT1/2 together increased ERK1/2 *S*-acylation; in fact, a decrease was observed (Figure S5E). These results once again suggest that the steady-state level *S*-acylation of ERK2 is tightly regulated.

Given the challenges of identifying the eraser via observations of steady-state ERK1/2 *S*-acylation, we generated myc-ERK2 tagged with TurboID, a biotin ligase that generates biotin–AMP to rapidly label proximal proteins and enable the mapping of the protein-protein interactions (Cho et al., 2020). After validating the expression and ligase activity of the constructs, we used this TurboID-tagged myc-ERK2 to probe its association with APT1 and APT2 (Figure S5F). Here, we observed labeling of and thus proximity to APT2, but not APT1 (Figure 5D, S5F). To further confirm this observation and establish the relevance of APT2 in ERK2’s dynamic *S*-acylation, we next carried out co-immunoprecipitation of APT2 with myc-ERK2 with and without EGF stimulus. This revealed a decrease in the interaction between ERK2 and APT2 upon the addition of EGF, an interaction that was also diminished upon treatment with PalmB (Figure 5E). In total, these observations further substantiate a model of dynamic ERK *S*-acylation – modulated by the interplay of several DHHCs and APT2 – in the regulation of its EGF-induced activity.

### ERK1/2 *S*-acylation is altered under conditions of metabolic stress

ERK1/2 signaling is not only responsive to growth factors, but also to cellular metabolism, with diet-derived molecules, such as glucose and long chain fatty acids, influencing enzyme activity and gene expression (Gehart et al., 2010). In fact, dysregulated ERK signaling, resulting from aberrant nutrient levels, has been implicated in pathogenesis of metabolic syndrome (Ozaki et al., 2016). While the mechanisms of this phenomenon are unknown, metabolite-dependent PTMs, such as *S*-acylation, present an intuitive, albeit unestablished, regulatory mechanism (Spinelli et al., 2018). Thus, we aimed to determine if changes in *S*-acylation of ERK1/2 could contribute to the changes in its activity *in cellulo* and *in vivo*. Incubation of HepG2 cells with a bolus of palmitate (500 μM), a condition known to induce metabolic stress and alter ERK1/2 activation, decreased ERK1/2 *S*-acylation levels, confirming the sensitivity of ERK1/2 *S*-acylation to metabolic stressors (Figure 6A).

**Figure 6.**
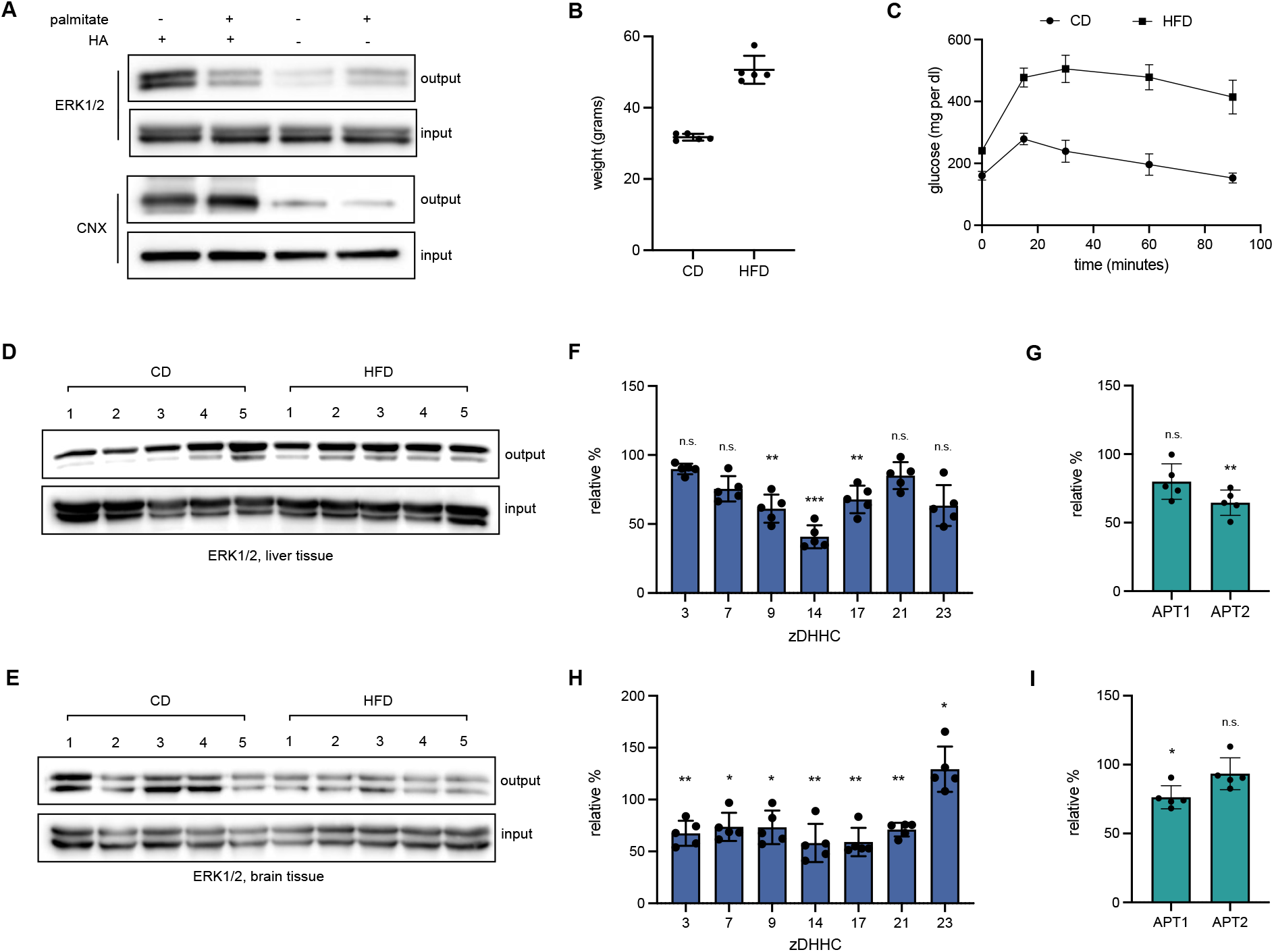
ERK1/2 *S*-acylation is responsive to metabolic stress *in cellulo* and *in vivo*. (**A**) ABE assay in HepG2 cells treated with palmitate (0 or 500 μM). (**B**) Plot of the weight of mice fed either a control diet (CD) or a high-fat diet (HFD). *mean ± std, n = 5*. (**C**) Glucose tolerance test (GTT) for mice fed either a control diet (CD) or a high-fat diet (HFD). *mean ± std, n = 5*. ABE assay carried on the liver (**D**) or brain (**E**) tissues of C57BL/6 fed either a control (CD) or high-fat (HFD) diet for 20 weeks. (**F**) RT-qPCR analysis of ERK DHHC writer transcript levels in the liver of HFD mice, relative to those of CD mice. (**G**) RT-qPCR analysis of APT transcript levels in the liver of HFD mice, relative to those of CD mice. (**H**) RT-qPCR analysis of ERK DHHC writer transcript levels in the brain of HFD mice, relative to those of CD mice. (I) RT-qPCR analysis of APT transcript levels in the brain of HFD mice, relative to those of CD mice. Statistical analyses performed with a two-tailed student’s t-test with equal variance (*n=5*). * = p < 0.5, ** = p < 0.005, and *** = p < 0.005.

We next assessed changes in ERK1/2 *S*-acylation in relevant tissues in a mouse model of diet-induced obesity. C57BL/6J were fed either a high-fat diet (HFD) rich in palmitate or a matched control diet (CD) for 18 weeks, and then evaluated for metabolic health (Figure 6B,C, S6) (Wang & Liao, 2012). After confirming the onset of metabolic syndrome in HFD mice, we probed for changes in ERK1/2 *S*-acylation in the liver, a metabolically active tissue in which ERK signaling is dysregulated in obesity, and the brain, a tissue with known sensitivity to circulating palmitate levels (Bi et al., 2013; Bost et al., 2005; Karmi et al., 2010). Here, we observed an increase in the *S*-acylation of ERK1/2, especially for ERK2, in the liver, while a decrease was observed in the brain (Figure 6D,E). Moreover, transcript abundance estimation via RT-qPCR indicated corresponding changes in the expression of ERK2 writers, with DHHCs 9, 14, and 17 and APT2 downregulated in the liver and DHHCs 3, 7, 9, 14, 17, 21, and 23 downregulated in the brain (Figure 6F-I). Intriguingly, the decrease in APT expression in the liver paralleled the increase in ERK S-acylation, while the decrease in DHHC expression in the brain liver paralleled the decrease in ERK S-acylation. These results suggest the enzyme-mediated changes in the S-acylation of ERK1/2 could contribute to the mechanism of dysregulated ERK1/2 signaling observed across tissues in metabolic syndrome.

## Discussion

As members of the conserved MAPK family, ERK1/2 contribute to essential cell processes and are associated with cell proliferation, cell growth, cell mobility, and cell survival (Roskoski, 2012). Targeting ERK1/2 activity is of therapeutic interest for a constellation of pathological conditions, including cancer, metabolic syndrome, neurological disease, and chronic inflammation (Lu & Malemud, 2019; Miao & Tian, 2020; Ozaki et al., 2016; Sun & Nan, 2017). Here, we introduce *S*-acylation as a novel, signal-responsive PTM for ERK1/2, in particular ERK2. The dynamics observed – i.e., increasing *S*-acylation with EGF treatment – is in accordance with our previously reported rapid decrease in APT activity upon EGF treatment and highlights the regulatory role of this PTM and the potential of APT activity to tune substrate lipidation levels (Beck et al., 2017; Kathayat et al., 2017; Qiu et al., 2018). In addition, when coupled with the observed specificity of the increase (EGF over insulin) and the previously described role of phosphorylation in regulating APT1 activity, this suggests an as-yet undescribed cellular conversation between the acylation regulatory machinery and canonical phosphorylation cascades (Sadeghi et al., 2018). Defining the nature of these upstream interactions represents a new and significant line of investigation.

Our experiments also highlight the relevance of *S*-acylation for ERK2 activity and function. Although mutation (activation/inactivation) of the canonical TEY motif did not disrupt the dynamics of ERK2 *S*-acylation, biochemical perturbation of a primary acylation site of ERK2, C254, resulted in an altered pattern of phosphorylation for both its activation loop TEY motif and serine residues. Given the observed changes in TEY motif phosphorylation, which is significant for ERK2 activation, we also assessed the effect of disrupting the cycle of ERK1/2 *S*-acylation on their transcriptional program. Here, we observed that chemical inhibition of both lipid installation and removal disrupted the activation timing and amplitude of targets like EGR1, cFos, and DUSP6 (Figure 4C). This suggests that an inhibitor targeting C254 of ERK2 or possibly C271 of ERK1 could be used to regulate their activity and downstream cellular events. The potential of cysteine-reactive chemical tools to moderate ERK1/2 is also highlighted by the observation that the decreased expression of ERK1/2 coincident with the mutagenesis of Cys233/216 can be reversed by the addition of either the proteasome inhibitor MG132 or the APT2 inhibitor ML-349, although further studies are needed to understand the intersection between *S*-acylation and ubiquitination-dependent degradation (Figure S6B). Finally, while the implications of serine phosphorylation for ERK2 are incompletely established, it has been postulated that a serine motif regulates ERK1/2 interactions with importin7 (Plotnikov et al., 2019). Intriguingly, importin7 is a lipid-binding protein, and an ERK nuclear translocation signal (NTS)-derived myristoylated phosphomimetic peptide has been shown to inhibit the ERK2-importin7 interaction, suggesting that ERK2 acylation could be part of a coincidence detection motif in this protein-protein interaction (Niphakis et al., 2015; Plotnikov et al., 2015). Alterations in the ERK/importin7 interaction and ensuing changes in the timing of nuclear import could also explain the altered transcriptional program timing (Figure 4C).

We additionally identified the enzymatic mediators of ERK1/2’s acylation and deacylation, with zDHHCs 7 and 21 emerging as two of several writers, and APT2, as a key eraser (Figure 5). Interestingly, traditional methods to identify writers and erasers, including both overexpression and knockdown of said proteins, failed to reveal DHHC and APT regulators of ERK *S*-acylation. This result emphasizes that the identification of molecular determinants for a substrate whose steady state *S*-acylation status is tightly regulated by multiple actors requires different experimental approaches. In the case of ERK2, chemical metabolic labeling enabled observation of changes in palmitate incorporation with DHHC overexpression, while co-immunoprecipitation validated DHHC/ERK2 interactions and the phosphorylation change assay, the role of DHHCs in ERK2 activation. The observation of dynamic interactions between ERK2 and the DHHCs/APT2 in the context of EGF stimulation is also noteworthy. While steady-state substrate-writer/eraser associations are well-established (e.g., STAT3/DHHC7 and Scribble/APT2), here, we find that interactions between ERK2 and DHHCs/APT2 are sensitive to EGF signaling (Hernandez et al., 2017; Zhang et al., 2020). This again hints at upstream crosstalk between dynamic phosphorylation and *S*-acylation, and, although the mechanisms that drive these interactions have yet to be elucidated, it also spotlights the importance of exploring dynamic interactions in changing cellular contexts for other *S*-acylated substrates.

In identifying ERK1/2 *S*-acylation writers, we also observed that certain DHHC family proteins, including 9, 23, 12, 15, and 20, appeared to affect the *S*-acylation of ERK1 more significantly than that of ERK2 (Figure 5A). These results suggest a possible ERK isoform-specific mode of regulation and highlights the potential of isoform-selective DHHC inhibitors to modulate ERK1 vs. ERK2 signaling. In addition, the efficacy of CMA as a chemical inhibitor of ERK1/2 *S*-acylation, in contrast to the well-established 2BP, emphasizes the potential utility of even new broad-spectrum DHHC family inhibitors. Also interesting was ERK2’s association with zDHHC5 (Figure S5C). This plasma-membrane localized PAT is unlikely a writer of ERK1/2 S-acylation, given its inability to increase fatty acid incorporation (Figure 5A) (Ohno et al., 2006). However, proximity proteomics experiments indicate that it is associated with Grb2, an adaptor protein that links the MAPK pathway to activated RTKs (Ke et al., 2021). These associations suggest it may have a role as a scaffolding protein in the EGFR signaling complex.

The observation of organ-specific changes in ERK1/2 *S*-acylation in a mouse model of metabolic syndrome underscores the responsiveness of this modification to the organismal environment and its potential as a regulatory mechanism (Figure 6). Given our data that suggest ERK1/2 acylation status regulates their activity and stability, this result strongly suggests that *S*-acylation could be a significant contributor to altered ERK1/2 signaling in metabolic syndrome. The alterations in expression of *S*-acylation writers and erasers suggest that the acylation machinery is sensitive to metabolic stress – an observation with implications not only for ERK1/2 but also for the panel of proteins whose activity is regulated by enzymatic *S*-acylation.

Overall, this works helps to address the still salient question for ERK1/2 – *quis regit ipsos regendos/who regulates the regulators?* – by adding a new element to the regulation of ERK1/2 and presenting a new therapeutic hypothesis for targeting ERK1/2-driven pathologies. More broadly, this work highlights the connection between dynamic lipidation and phosphorylation, a connection hinted by our previous work showing that APT activity is responsive to growth factor stimulation and lipid stress and by the observation that APT1 activity is regulated by phosphorylation (Kathayat et al., 2017; Qiu et al., 2018; Sadeghi et al., 2018). It also underscores the importance of timing and cell state in identifying the molecular determinants and significance of *S*-acylation for a substrate, especially given the observation of dynamic interactions between ERK2 and APT2/DHHCs. Further exploration of the mechanisms that regulate the dynamic activity and interactions of the acylation writer/eraser proteins will broaden our perspective on the regulatory machinery of the cell in both physiological and pathological contexts. This work and the approaches described herein grow our understanding of dynamic protein *S*-acylation and provide a roadmap to further deconvolute the role of dynamic *S*-acylation in regulating signal transduction.

---

**Figure S1.**
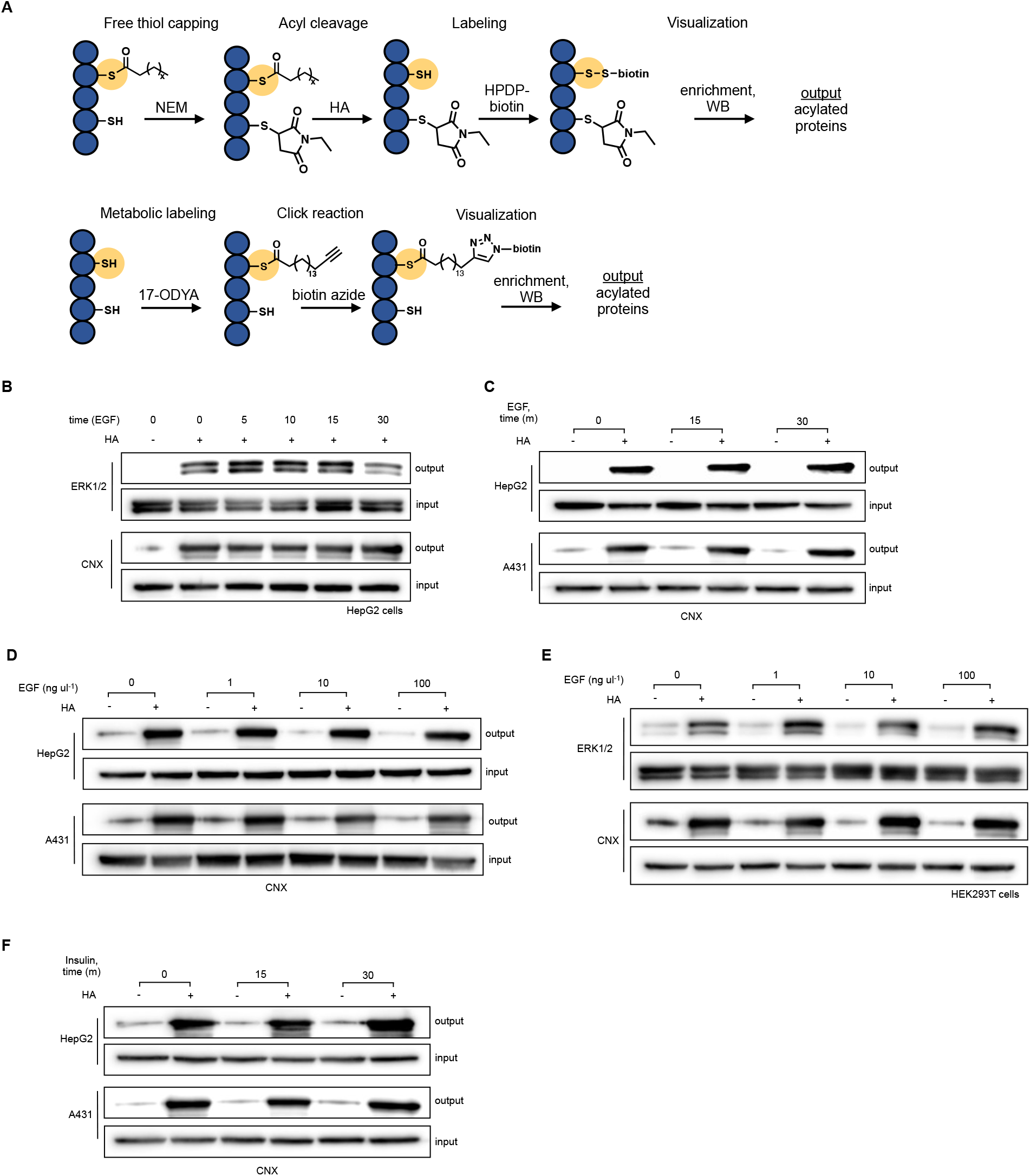
The extracellular signaling regulated kinases (ERK1/2) are dynamically palmitoylated. (**A**) Schematic representations of two key assays, acyl biotin exchange (ABE, top) and metabolic labeling (bottom) used to visualize protein *S*-acylation. (**B**) ABE assay carried out in HepG2 following stimulation with EGF (1 ng mL^−1^) for 0, 5, 10, 15, and 30 minutes. (**C**) Western blots for calnexin (CNX), used as both a loading and assay control for 2A. (**D**) Western blots for calnexin (CNX), used as both a loading and assay control for 2B. (**E**) ABE assay carried out in HEK293T cells following stimulation with 0, 1, 10, and 100 ng mL^−1^ EGF for 15 minutes. (**F**) Western blots for calnexin (CNX), used as both a loading and assay control for 2C.

**Figure S2.**
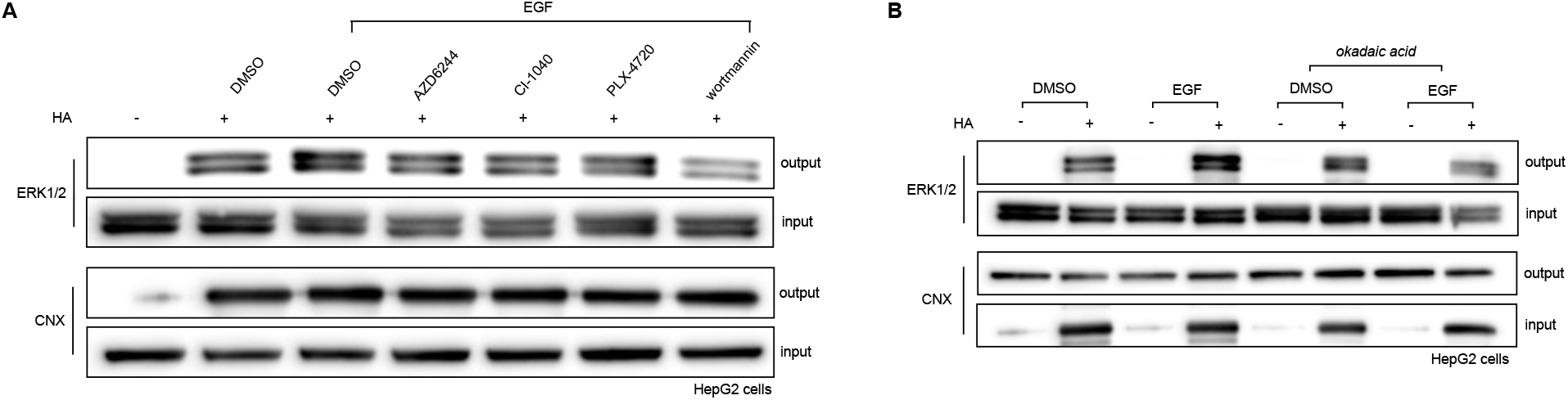
Dynamic S-acylation of ERK1/2 relies on pathway activation. (**A**) ABE assay on HepG2 cells treated with DMSO or MAPK and PI3K pathway inhibitors, followed by EGF treatment (15 minutes, 1 ng mL^−1^). (**B**) ABE assay carried out on HepG2 cells treated with DMSO or okadaic acid, followed by EGF treatment (15 minutes, 1 ng mL^−1^).

**Figure S3.**
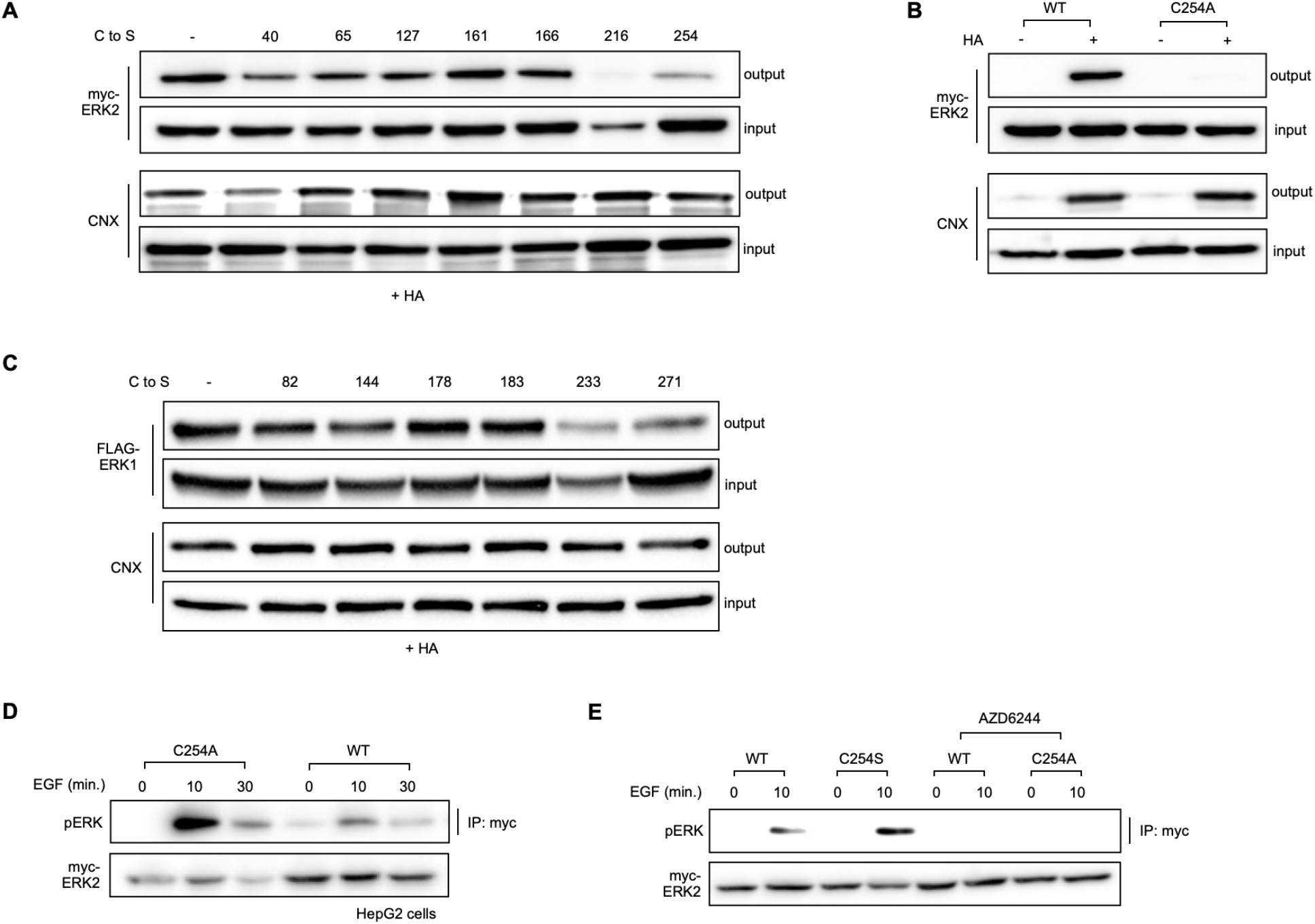
Identification of ERK1/2 acylation sites to probe for effects on phosphorylation. (**A**) ABE carried out on HEK293T cells overexpressing a panel of myc-ERK2 C to S mutants. (**B**) ABE of HEK293T cells expressing either WT or C254A myc-ERK2. (**C**) ABE carried out on HEK293T cells overexpressing a panel of 3xFLAG-ERK1 C to S mutants. (**D**) Immunoprecipitation of WT or palmitoylation-deficient myc-ERK2 in HepG2 cells stimulated with EGF (0, 10, and 30 minutes, 1 ng mL^−1^). (**E**) Immunoprecipitation of WT or palmitoylation-deficient myc-ERK2 in HEK293T cells stimulated with EGF (0, and 10 minutes, 1 ng mL^−1^), with or without MEK inhibitor AZD6244.

**Figure S4.**
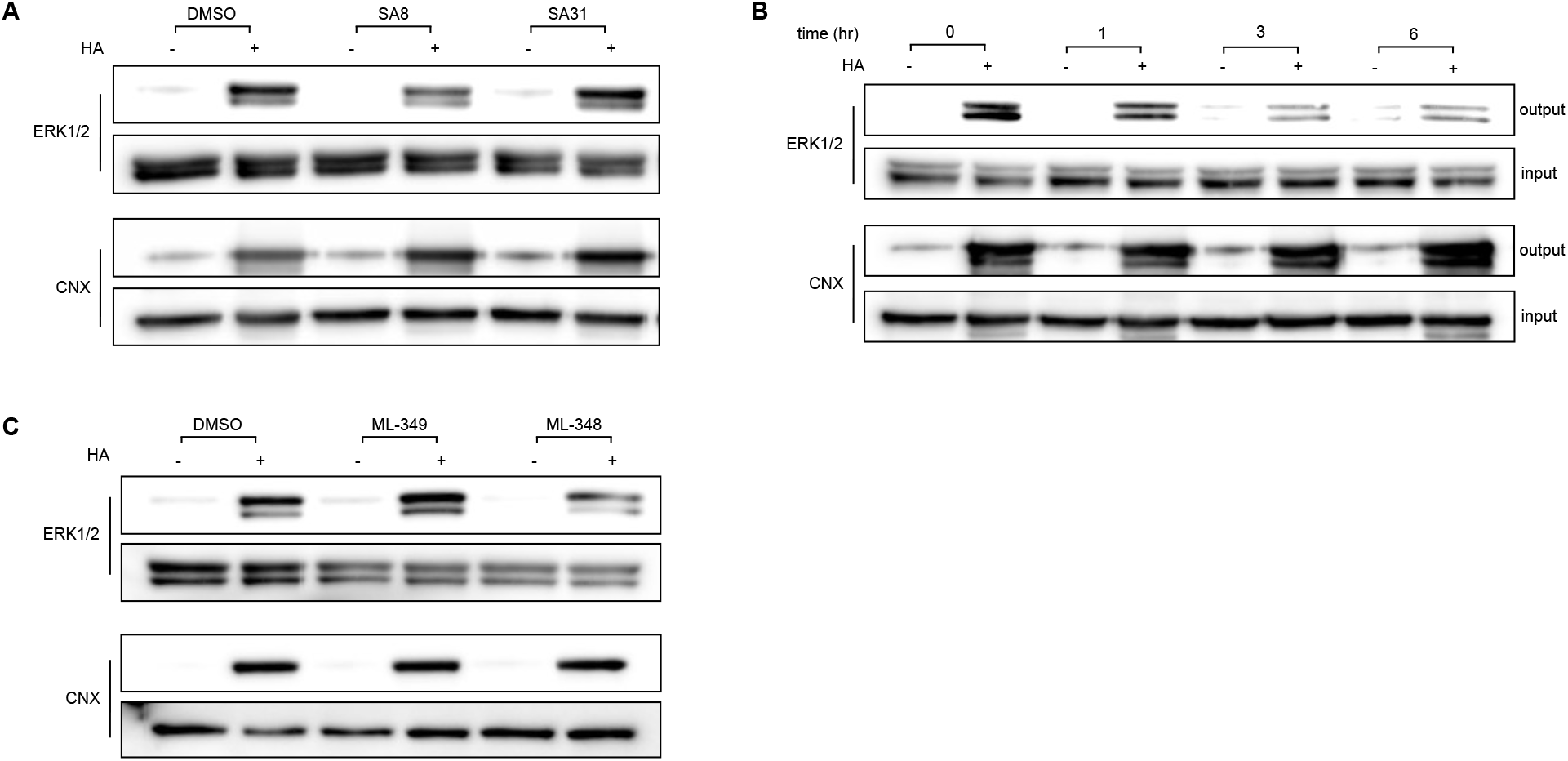
Chemical inhibition of ERK1/2 *S*-acylation. (**A**) ABE of HepG2 cells treated with DMSO, CMA (20 μM, 3 hours), or 14 (20 μM, 3 hours), an inactive analogue of CMA. (**B**) Timecourse ABE (0, 1, 3, and 6 hours) of HepG2 cells treated with CMA (20 μM). (**C**) ABE of HepG2 cells treated with DMSO, ML-348 (20 μM, 3 hours), or ML-349 (20 μM, 3 hours).

**Figure S5.**
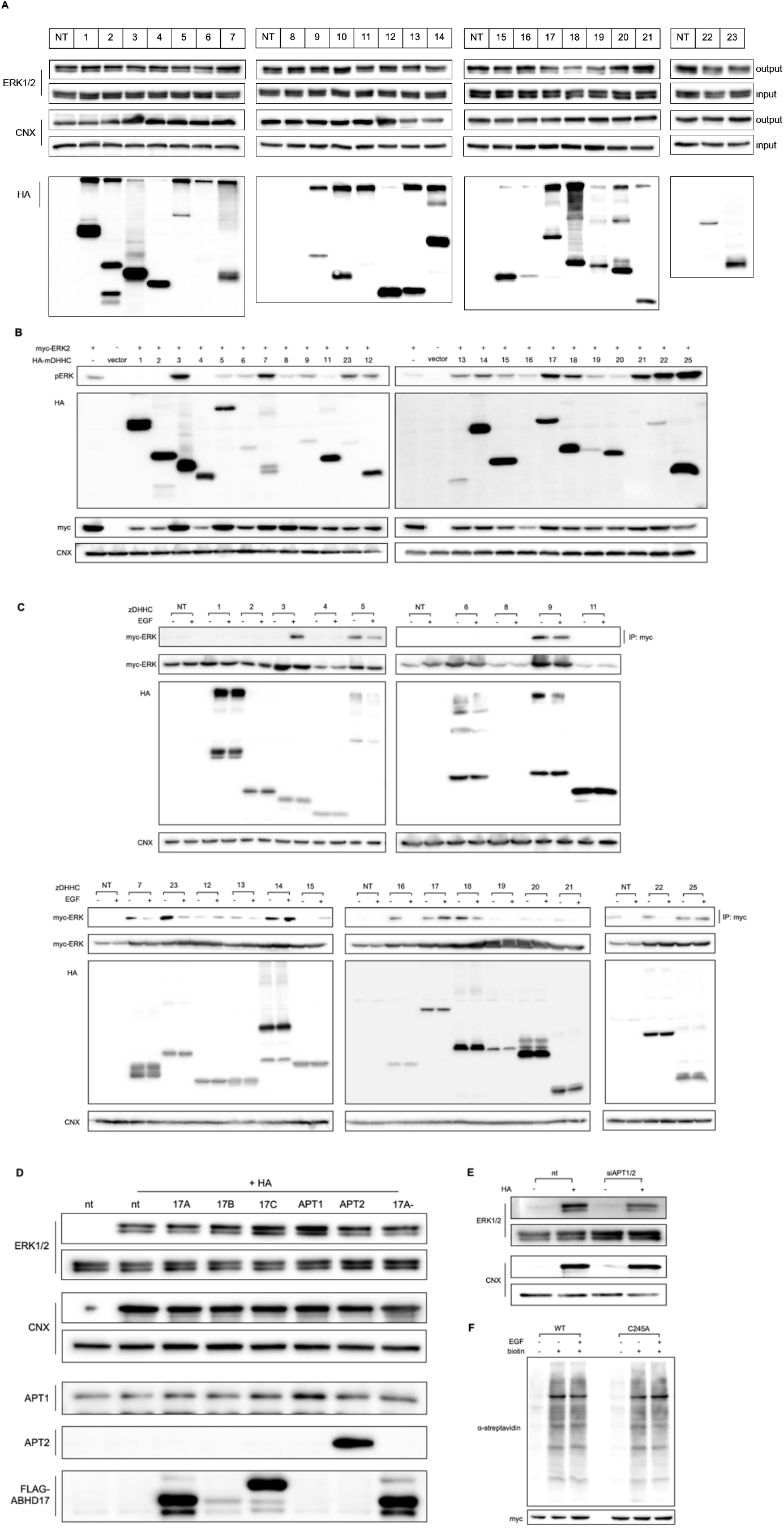
Identification of the writer and eraser enzymes mediating ERK2 *S*-acylation. (**A**) ABE of HEK293T cells overexpressing a panel of murine DHHC family proteins. (**B**) Analysis of phospho-ERK(Thr185/Tyr187) in HEK293T cells overexpressing a panel of murine DHHC proteins, following EGF stimulation (10 minutes, 1 ng mL^−1^). (**C**) Co-immunoprecipitation of myc-ERK2 with HA-tagged DHHC proteins in HEK293T cells with or without EGF stimulation (10 minutes, 1 ng mL^−1^). Co-immunoprecipitated proteins were visualized via Western blotting for myc-ERK2. (**D**) ABE of HEK293T cells overexpressing a panel of known APT enzymes. (**E**) ABE in HEK293T cells following siRNA-mediated knockdown of APT1 and APT2. (**F**) Validation of TurboID-tagged ERK2 constructs, with α-streptavidin staining indicating the activity of the biotin ligase and α-myc blotting indicating the expression levels of the constructs.

**Figure S6.**
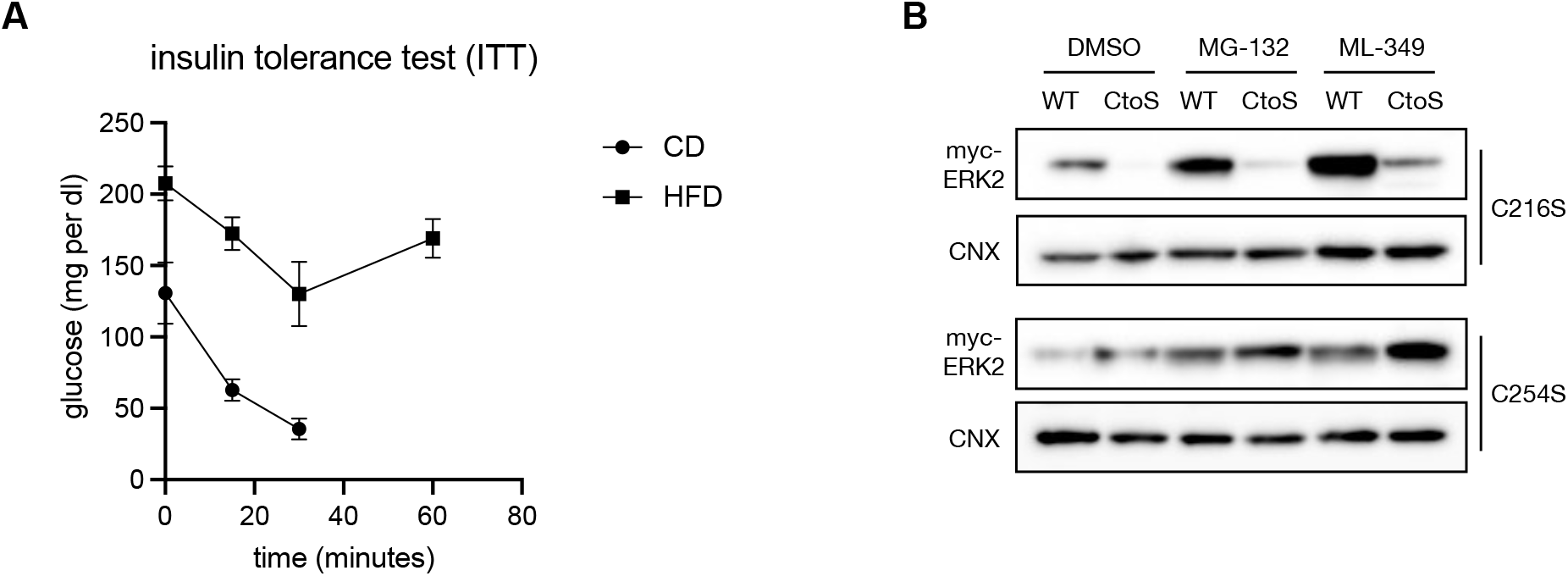
Characterization of DIO in C57BL/6J mice and characterization of ERK2 expression *in cellulo*. (**A**) Insulin tolerance test (ITT) for mice fed either a control diet (CD) or a high-fat diet (HFD). *mean ± std, n = 5*. (**B**) Overexpression of WT and C254S or C216S ERK2 in HEK293T cells treated with either the proteasome inhibitor MG-132 or the APT2 inhibitor ML-349. CNX was used as a loading control.

## Materials and Methods

Sources for common reagents are noted in relevant sections below. Short interfering RNAs (siRNAs) targeting human APT1/LYPLA1 (SI03246586) and human APT2/LYPLA2 (SI04269041), as well as nontargeting (NT) control siRNA (SI03650325), were purchased from Qiagen. Chemical inhibitors used in this study were sourced as follows: 2-bromopalmitate (Sigma); Palmostatin B (EMD Millipore); ML-348 (Tocris); and ML-349 (Tocris)

### Plasmid cloning

All plasmids were constructed by Gibson Assembly from PCR products generated using Q5 Hot Start DNA Polymerase (New England Biolabs) or Phusion Polymerase (generated in-house). Plasmids for human ERK1 (Plasmid #49328), human ERK2 (Plasmid #82145), and TurboID (Plasmid #124646) were obtained from Addgene and subsequently cloned into the d0 backbone. All newly constructed plasmids were sequence-verified by the University of Chicago Comprehensive Cancer Center DNA Sequencing and Genotyping Facility and are available on request.

### Cell culture

Cells were plated and maintained in growth media (DMEM GlutaMAX for HEK293T and A431cells; EMEM for HepG2 cells) supplemented with 10% FBS and 1% penicillin/streptomycin at 37°C and 5% CO_2_. For all experiments, cells had undergone fewer than 18 passages. Transfections were conducted using PEI (Sigma) or Lipofectamine 3000 (Invitrogen) in accordance with manufacturer protocols.

### Western blotting

After SDS–PAGE, proteins were transferred onto methanol-preactivated Immobilon-P PVDF membranes (pore size 0.45 μm; Millipore) using a semi-dry transfer cell or a wet transfer tank (Bio-Rad). After transfer, membranes were treated in accordance with standard Western blotting procedures, using a solution of 3% BSA (ThermoFisher) in TBST (20 mM Tris, pH 7.5, 150 mM NaCl, 0.1% Tween-20) wash buffer. Membranes were visualized using SuperSignal West Pico PLUS chemiluminescent substrate (ThermoFisher) and recorded on a chemiluminescent western blot imaging system (Azure Biosystems C300). All antibodies used can be found in Table 1.

**Table 1.**
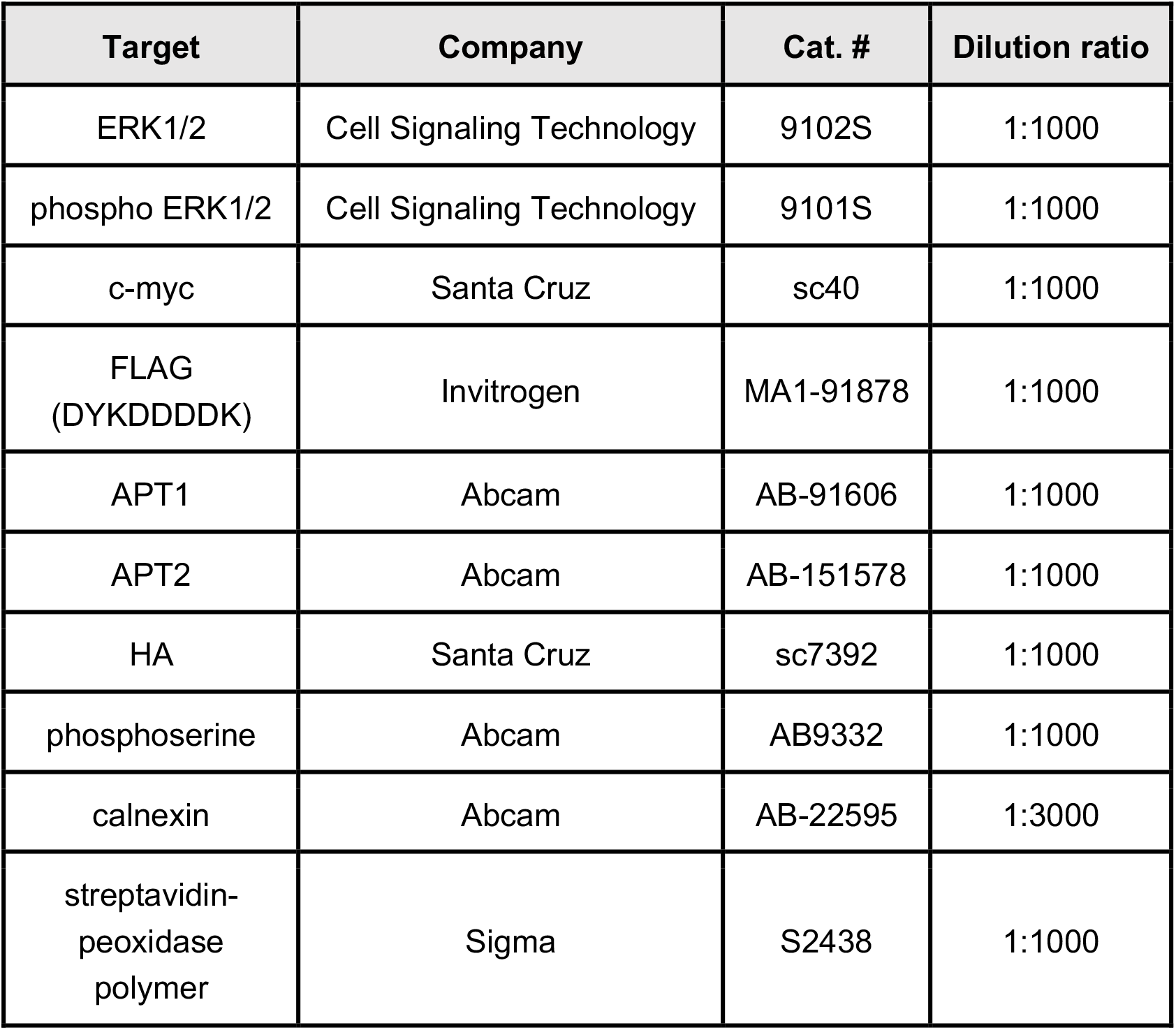
Antibodies

| Target | Company | Cat. # | Dilution ratio |
| --- | --- | --- | --- |
| ERK1/2 | Cell Signaling Technology | 9102S | 1:1000 |
| phospho ERK1/2 | Cell Signaling Technology | 9101S | 1:1000 |
| c-myc | Santa Cruz | sc40 | 1:1000 |
| FLAG<br>(DYKDDDDK) | Invitrogen | MA1-91878 | 1:1000 |
| APT1 | Abcam | AB-91606 | 1:1000 |
| APT2 | Abcam | AB-151578 | 1:1000 |
| HA | Santa Cruz | sc7392 | 1:1000 |
| phosphoserine | Abcam | AB9332 | 1:1000 |
| calnexin | Abcam | AB-22595 | 1:3000 |
| streptavidin-<br>peoxidase<br>polymer | Sigma | S2438 | 1:1000 |

### Acyl biotin exchange (ABE)

#### All volumes and reagent amounts are representative for an experiment using a 10 cm plate

Following treatment as required, cells were washed twice with cold DPBS and lysed with 1-2 mL of RIPA lysis buffer (50 mM Tris, 150 mM NaCl, 0.5% deoxycholate, 0.1% SDS, 1.0% Triton X-100, pH 7.4) supplemented with a protease inhibitor cocktail and 50 mM *N*-ethylmaleimide (NEM) (Acros). After end-over-end rotation (4 °C, 3 hours), lysates were centrifuged at 13,000*g* (4 °C, 20 minutes), and the supernatant was collected and subjected to acetone precipitation. The resulting pellet was dissolved by sonication in resuspension buffer (4% SDS, 50 mM HEPES, 150 mM NaCl, 5 mM EDTA, pH 7.4; 100 μL per mg of protein) containing 50 mM NEM. This protein solution was rotated end-over-end (25 °C, 2 hours) and then subjected to chloroform–methanol precipitation (2x) to remove excess NEM. The resulting protein pellet was dissolved in 160 μL of resuspension buffer via sonication. The protein sample was divided into two equal parts for ±hydroxylamine (HA) (Combi-Blocks) treatment. Each sample was treated with 320 μL of either -HA Triton buffer (0.2% Triton X-100, 50 mM HEPES, 150 mM NaCl, 5 mM EDTA, pH 7.4) or +HA buffer (Triton buffer +0.7 M HA, pH ~7.2). After shaking incubation (25 °C, 1 hour), proteins were precipitated by chloroform–methanol precipitation to remove HA. Protein pellets were resuspended by sonication in 80 μL of resuspension buffer containing 10 μM EZ-Link HPDP-Biotin (ThermoFisher), diluted with 320 μL of Triton buffer containing 10 μM HPDP-Biotin, and incubated with shaking (25 °C, 2 hours). Excess biotin was removed with two sequential chloroform-methanol precipitations. Finally, protein pellets were dissolved in 40 μL resuspension buffer via sonication, and the solution volume was brought to 400 μL with Triton buffer. Protein concentration was then measured using the BCA assay (ThermoFisher). Protein (~30-40 μg, 10%) was removed to serve as an expression control (‘input’). The remaining solution was diluted with Triton buffer, and streptavidin–agarose beads (50 μL of slurry per 1 mg of protein) were added to each sample, which was then incubated via end-over-end rotation (4 °C, 12 hours). Beads were then washed (3 × 1 mL) with wash buffer (Triton buffer + 0.1% SDS) to remove unbound protein. Bound proteins (‘output’) were eluted by boiling the beads (95 °C, 10 minutes) with 1x Laemelli sample buffer (Alfa Aesar) containing 30 mM DTT. The protein was resolved on 8-12% SDS–PAGE gels and subjected to Western blotting in accordance with the protocol outlined above.

For ABE performed on mouse tissue samples, organs were isolated from C57BL/6J mice, washed with Hanks’ balanced salt solution, and flash-frozen in liquid nitrogen. Following mechanical homogenization, samples were lysed in 2–5 ml (depending on organ mass) of HEPES lysis buffer (150 mM NaCl, 40 mM HEPES, 0.5% Triton, 3 mM EDTA, pH 7.4 with protease inhibitors and 1 mM PMSF) containing 50 mM NEM, vortexed vigorously and rotated end-over-end (4 °C, 8 hours). After centrifugation at 15,000*g*, the supernatant was removed and subjected to sequential acetone and chloroform–methanol precipitations to yield dry protein. The resulting protein pellet was then processed as described above.

To visualize phosphorylated and acylated ERK1/2, phosphatase inhibitor cocktails (Santa Cruz, sc-45044 and sc-45045) were added to the lysis buffer and to subsequent protein solutions throughout the assay.

For ABEs following chemical inhibitor treatment (CMA, PalmB, ML-348, and ML-349), cells were treated at the stated concentrations and times in media containing 0.1% FBS (DMEM, HEK293T cells) or 1% FBS (EMEM, HepG2 cells).

For ABEs following mDHHC library overexpression HEK293T cells first were transfected with HA-tagged mDHHC plasmids (11 μg DNA per 10 cm dish) at 40-60% confluency. At 24 hours post transfection, HEK293T cells were washed with DPBS and lysed with 1-2 mL of RIPA lysis buffer (50 mM Tris, 150 mM NaCl, 0.5% deoxycholate, 0.1% SDS, 1.0% Triton X-100, pH 7.4) supplemented with a protease inhibitor cocktail and 50 mM *N*-ethylmaleimide (NEM) (Acros). The resulting lysate was then processed as described above.

For ABEs following APT knockdown in a 10 cm dish, HEK293T cells at 40-60% confluency were transfected with 10 pmol of siRNA targeting either APT1 or APT2, or a NT siRNA control. At 24 hours post transfection, HEK293T cells were washed with DPBS and lysed with 1-2 mL of RIPA lysis buffer (50 mM Tris, 150 mM NaCl, 0.5% deoxycholate, 0.1% SDS, 1.0% Triton X-100, pH 7.4) supplemented with a protease inhibitor cocktail and 50 mM *N*-ethylmaleimide (NEM) (Acros). The resulting lysate was then processed as described above.

### Metabolic Labeling

#### All volumes and reagent amounts are representative for an experiment using 2 wells of 6 well plate

At 40-60% confluency, HEK293T cells were transfected with HA-tagged mDHHC plasmids (2 μg DNA/well). At 24 hours post transfection, HEK293T cells were washed with DPBS and treated with 50 μM 17-ODYA absorbed on 5% BSA in DMEM GlutaMAX supplemented with 5% charcoal filtered FBS (6 hours). Following treatment, cells were washed once in DPBS, lysed with 280 μL HEPES Lysis buffer (150 mM NaCl, 50 mM HEPES, 0.2% SDS, 1% Triton-100, pH 7.4, supplemented with an EDTA-free protease inhibitor cocktail and 1mM PMSF), and subjected to end-over-end rotation (4 °C, 12 hours). The cell debris was pelleted by centrifugation at 13000*g* (4 °C, 20 minutes) and the supernatant collected. A master mix (24 μL, per sample, 6 μL each of 5 mM Biotin-Azide, 5 mM TBTA, 50 mM CuSO_4_ and 5 mM TCEP) was made and added for a 300μL “click” reaction. The resulting solution was incubated (r.t., 1 hour) with shaking. 10 μL of 100 mM EDTA was added to quench the “click” reaction and proteins were precipitated by chloroform-methanol precipitation (2x). The resulting protein pellet was dissolved in 40 μL of the resuspension buffer via sonication and the volume was brought to 1600 μL with Triton buffer. Undissolved material was removed by centrifugation at 13,000 g (4 °C, 5 minutes). Two 40 μL protein samples were transferred to fresh tubes (‘input’ and expression validation). 25 uL of streptavidin-agarose bead slurry was added to 1400uL of the remaining protein solution, and then samples were incubated with end-over-end rotation (4 °C, 12 hours). Unbound proteins were removed by washing (6x 1 mL) with wash buffer. Bound proteins were eluted by boiling the beads (95 °C, 10 minutes) with 1x Laemelli sample buffer containing 20-30 mM DTT. The protein was resolved on 8-12% SDS-PAGE gels and subjected to Western blotting protocol using the protocol described above.

### Immunoprecipitations

For immunoprecipitation and co-immunoprecipitation experiments, following stated transfections and treatments, cells were lysed in SDS-free RIPA (150 mM Tris, 150 mM NaCl, 0.5% deoxycholate, 1.0% Triton X-100, pH 7.4) supplemented with PMSF and a protease inhibitor cocktail on ice for 10 minutes, followed by centrifugation at 13,000*g* (4 °C, 20 min). Protein concentration was then measured using the BCA assay (ThermoFisher), and protein (20 μg, ‘input’) was removed and lysate was added to Protein G Dynabeads (Invitrogen: 10003D) pre-incubated with the appropriate antibody. After 12 hours of enrichment, beads were washed (3x, SDS-free RIPA) and total protein eluted with 50 mM glycine (pH 2.8) and brought to neutral with 1 μL of 1 M Tris-base, ‘output.’ Both input and output protein were subjected to separation by SDS-PAGE and Western blotting in accordance with described procedures.

### RT-qPCR

For *in cellulo* experiments, HepG2 cells were starved in low-serum media (1% EMEM) for 16 hours, and then treated first with DMSO, CMA (20 μM, 3 hours), or PalmB (20 μM, 3 hours) and then with EGF (1 ng mL^−1^) for 0, 30 or 60 minutes. Total RNA was harvested and isolated using the RNeasy Mini Kit (QIAGEN) and RNA clean & concentration (Zymo). After RNA isolation, RNA was reverse transcribed using the PrimeScript RT Reagent Kit (TaKaRa). All qPCR reactions were run at 20 μL volumes with three biological replicates using PowerUp SYBR Green Master Mix on a QuantStudio 3 Real-Time PCR System (Thermo). Relative expression levels for target genes were calculated using the 2^−ΔΔ^Ct method, with gene expression first compared to the housekeeping control gene (β2M) cycle threshold (Ct) value and then to the DMSO-treated samples. For mouse experiments, tissue samples were homogenized in Trizol and subjected to phase separation and precipitation to isolate the total RNA. After RNA isolation, RT-qPCR steps were carried out as described above. Relative expression levels for target genes were calculated using the 2^−ΔΔ^Ct method, with gene expression in DIO mice first compared to the housekeeping control gene (β2M) cycle threshold (Ct) value and then to the DIO-control samples. All qPCR primers can be found in Table 2.

**Table 2.**
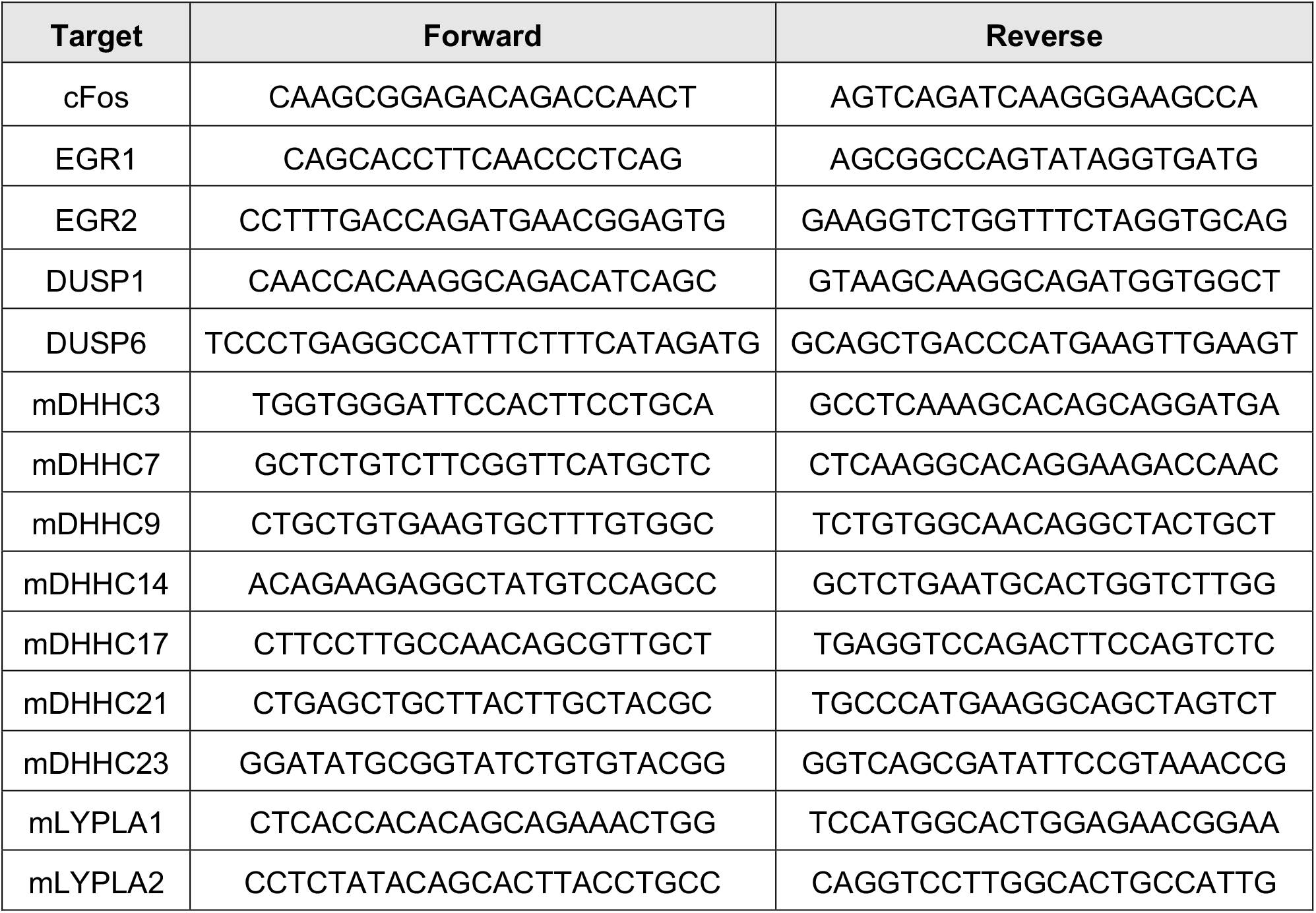
RT-qPCR primers

### TurboID

At 40-60% confluency, HEK293T cells were transfected with ERK2(WT)-TurboID or ERK2(Mutant)-TurboID (10 μg DNA/10 cm dish). At 24 hours post transfection, HEK293T cells were starved in 0.1% FBS DMEM media for 12 hours. Then the cells were treated with 100 μM Biotin (Sigma) in 0.1% FBS DMEM with or without 1 ng/mL EGF for 12 minutes. The cells were washed three times with DPBS and lysed in SDS-free RIPA buffer supplemented with PMSF and protease inhibitors on ice for 30 minutes, followed by centrifuge at 13,000*g* (4 °C, 20 min.) Protein (20 μg, ‘input’) was removed and 750ug lysate was added with streptavidin–agarose beads (30 μL of slurry), which was then incubated via end-over-end rotation (4 °C, 12 hours). Beads were then washed (3 × 1 mL) with wash buffer (0.1% SDS, 0.2% Triton X-100, 50 mM HEPES, 150 mM NaCl, 5 mM EDTA, pH 7.4) to remove unbound protein. Bound proteins (‘output’) were eluted by boiling the beads for 10 min at 95 °C with 1.5 × Laemelli sample buffer (Alfa Aesar) containing 30 mM DTT. The protein was resolved on 8-12% SDS–PAGE gels and subjected to Western blotting in accordance with the protocol outlined above.

### Evaluation of mouse metabolic health

The use of vertebrate animals (*Mus musculus*, mouse) in the laboratory of B. Dickinson has been approved by the Institutional Animal Care and Use Committee under the Animal Care and Use protocol no. 72531, ‘Mouse models of lipid signaling regulation’. DIO (Stock No: 380050) or DIO control mice (Stock No: 380056 (male, 5x each) were purchased from Jackson Labs at 12 weeks of age and then maintained on either the high fat diet (D12492, 60 kcal% fat) or the control diet (D12450B, 10 kcal% fat) (Research Diets, Inc.) for an additional 6 weeks. To confirm the manifestation of metabolic syndrome, insulin tolerance tests (ITT) and glucose tolerance tests (GTT) were carried out in accordance with previously reported protocols (Wang & Liao, 2012).

## Acknowledgements

This research was supported by the University of Chicago, the National Institute of General Medical Sciences of the National Institutes of Health NIH (R35 GM119840 to B.C.D.), and the National Institute of Diabetes and Digestive and Kidney Diseases of the National Institutes of Health (F30 DK125088 to S-A.A.). We thank the Fukata lab for providing us with the HA-tagged mDHHC plasmids. We would also like to thank Dr. S. Ahmadiantehrani and Prof. Marsha Rosner for assistance in preparing this manuscript, and Dr. M. Beck for early guidance in this project.

## References

Adibekian, A., Martin, B. R., Chang, J. W., Hsu, K. L., Tsuboi, K., Bachovchin, D. A., Speers, A. E., Brown, S. J., Spicer, T., Fernandez-Vega, V., Ferguson, J., Cravatt, B. F., Hodder, P., & Rosen, H. (2010a). Characterization of a Selective, Reversible Inhibitor of Lysophospholipase 1 (LYPLA1). In Probe Reports from the NIH Molecular Libraries Program. https://www.ncbi.nlm.nih.gov/pubmed/24624465

Adibekian, A., Martin, B. R., Chang, J. W., Hsu, K. L., Tsuboi, K., Bachovchin, D. A., Speers, A. E., Brown, S. J., Spicer, T., Fernandez-Vega, V., Ferguson, J., Cravatt, B. F., Hodder, P., & Rosen, H. (2010b). Characterization of a Selective, Reversible Inhibitor of Lysophospholipase 2 (LYPLA2). In Probe Reports from the NIH Molecular Libraries Program. https://www.ncbi.nlm.nih.gov/pubmed/24624468

Azizi, S.-A., Lan, T., Delalande, C., Kathayat, R. S., Banales Mejia, F., Qin, A., Brookes, N., Sandoval, P. J., & Dickinson, B. C. (2021). Development of an Acrylamide-Based Inhibitor of Protein S-Acylation. ACS Chemical Biology. https://doi.org/10.1021/acschembio.1c00405

Azizi, S. A., Kathayat, R. S., & Dickinson, B. C. (2019). Activity-Based Sensing of S-Depalmitoylases: Chemical Technologies and Biological Discovery. Acc Chem Res, 52(11), 3029–3038. https://doi.org/10.1021/acs.accounts.9b00354

Beck, M. W., Kathayat, R. S., Cham, C. M., Chang, E. B., & Dickinson, B. C. (2017). Michael addition-based probes for ratiometric fluorescence imaging of protein S-depalmitoylases in live cells and tissues. Chem Sci, 8(11), 7588–7592. https://doi.org/10.1039/c7sc02805a

Berti, D. A., & Seger, R. (2017). The Nuclear Translocation of ERK. Methods Mol Biol, 1487, 175–194. https://doi.org/10.1007/978-1-4939-6424-6_13

Bi, L., Chiang, J. Y., Ding, W. X., Dunn, W., Roberts, B., & Li, T. (2013). Saturated fatty acids activate ERK signaling to downregulate hepatic sortilin 1 in obese and diabetic mice. J Lipid Res, 54(10), 2754–2762. https://doi.org/10.1194/jlr.M039347

Blaustein, M., Piegari, E., Martinez Calejman, C., Vila, A., Amante, A., Manese, M. V., Zeida, A., Abrami, L., Veggetti, M., Guertin, D. A., van der Goot, F. G., Corvi, M. M., & Colman-Lerner, A. (2021). Akt Is S-Palmitoylated: A New Layer of Regulation for Akt. Front Cell Dev Biol, 9, 626404. https://doi.org/10.3389/fcell.2021.626404

Bolognesi, B., & Lehner, B. (2018). Reaching the limit. Elife, 7. https://doi.org/10.7554/eLife.39804

Bost, F., Aouadi, M., Caron, L., Even, P., Belmonte, N., Prot, M., Dani, C., Hofman, P., Pages, G., Pouyssegur, J., Le Marchand-Brustel, Y., & Binetruy, B. (2005). The extracellular signal-regulated kinase isoform ERK1 is specifically required for in vitro and in vivo adipogenesis. Diabetes, 54(2), 402–411. https://doi.org/10.2337/diabetes.54.2.402

Busca, R., Pouyssegur, J., & Lenormand, P. (2016). ERK1 and ERK2 Map Kinases: Specific Roles or Functional Redundancy? Front Cell Dev Biol, 4, 53. https://doi.org/10.3389/fcell.2016.00053

Cao, Y., Qiu, T., Kathayat, R. S., Azizi, S. A., Thorne, A. K., Ahn, D., Fukata, Y., Fukata, M., Rice, P. A., & Dickinson, B. C. (2019). ABHD10 is an S-depalmitoylase affecting redox homeostasis through peroxiredoxin-5. Nat Chem Biol, 15(12), 1232–1240. https://doi.org/10.1038/s41589-019-0399-y

Cargnello, M., & Roux, P. P. (2011). Activation and function of the MAPKs and their substrates, the MAPK-activated protein kinases. Microbiol Mol Biol Rev, 75(1), 50–83. https://doi.org/10.1128/MMBR.00031-10

Chamberlain, L. H., & Shipston, M. J. (2015). The physiology of protein S-acylation. Physiol Rev, 95(2), 341–376. https://doi.org/10.1152/physrev.00032.2014

Chang, F., Steelman, L. S., Lee, J. T., Shelton, J. G., Navolanic, P. M., Blalock, W. L., Franklin, R. A., & McCubrey, J. A. (2003). Signal transduction mediated by the Ras/Raf/MEK/ERK pathway from cytokine receptors to transcription factors: potential targeting for therapeutic intervention. Leukemia, 17(7), 1263–1293. https://doi.org/10.1038/sj.leu.2402945

Charron, G., Zhang, M. M., Yount, J. S., Wilson, J., Raghavan, A. S., Shamir, E., & Hang, H. C. (2009). Robust fluorescent detection of protein fatty-acylation with chemical reporters. J Am Chem Soc, 131(13), 4967–4975. https://doi.org/10.1021/ja810122f

Chen, S., Zhu, B., Yin, C., Liu, W., Han, C., Chen, B., Liu, T., Li, X., Chen, X., Li, C., Hu, L., Zhou, J., Xu, Z. X., Gao, X., Wu, X., Goding, C. R., & Cui, R. (2017). Palmitoylation-dependent activation of MC1R prevents melanomagenesis. Nature, 549(7672), 399–403. https://doi.org/10.1038/nature23887

Cho, K. F., Branon, T. C., Udeshi, N. D., Myers, S. A., Carr, S. A., & Ting, A. Y. (2020). Proximity labeling in mammalian cells with TurboID and split-TurboID. Nat Protoc, 15(12), 3971–3999. https://doi.org/10.1038/s41596-020-0399-0

Chuderland, D., Konson, A., & Seger, R. (2008). Identification and characterization of a general nuclear translocation signal in signaling proteins. Mol Cell, 31(6), 850–861. https://doi.org/10.1016/j.molcel.2008.08.007

Cohen, P., Klumpp, S., & Schelling, D. L. (1989). An improved procedure for identifying and quantitating protein phosphatases in mammalian tissues. FEBS Lett, 250(2), 596–600. https://doi.org/10.1016/0014-5793(89)80803-8

Dekker, F. J., Rocks, O., Vartak, N., Menninger, S., Hedberg, C., Balamurugan, R., Wetzel, S., Renner, S., Gerauer, M., Scholermann, B., Rusch, M., Kramer, J. W., Rauh, D., Coates, G. W., Brunsveld, L., Bastiaens, P. I., & Waldmann, H. (2010). Small-molecule inhibition of APT1 affects Ras localization and signaling. Nat Chem Biol, 6(6), 449–456. https://doi.org/10.1038/nchembio.362

Drisdel, R. C., & Green, W. N. (2004). Labeling and quantifying sites of protein palmitoylation. Biotechniques, 36(2), 276–285. https://doi.org/10.2144/04362RR02

Ebisuya, M., Kondoh, K., & Nishida, E. (2005). The duration, magnitude and compartmentalization of ERK MAP kinase activity: mechanisms for providing signaling specificity. J Cell Sci, 118(Pt 14), 2997–3002. https://doi.org/10.1242/jcs.02505

Feng, X., Sun, T., Bei, Y., Ding, S., Zheng, W., Lu, Y., & Shen, P. (2013). S-nitrosylation of ERK inhibits ERK phosphorylation and induces apoptosis. Sci Rep, 3, 1814. https://doi.org/10.1038/srep01814

Fukata, Y., Iwanaga, T., & Fukata, M. (2006). Systematic screening for palmitoyl transferase activity of the DHHC protein family in mammalian cells. Methods, 40(2), 177–182. https://doi.org/10.1016/j.ymeth.2006.05.015

Gehart, H., Kumpf, S., Ittner, A., & Ricci, R. (2010). MAPK signalling in cellular metabolism: stress or wellness? EMBO Rep, 11(11), 834–840. https://doi.org/10.1038/embor.2010.160

Hernandez, J. L., Davda, D., Cheung See Kit, M., Majmudar, J. D., Won, S. J., Gang, M., Pasupuleti, S. C., Choi, A. I., Bartkowiak, C. M., & Martin, B. R. (2017). APT2 Inhibition Restores Scribble Localization and S-Palmitoylation in Snail-Transformed Cells. Cell Chem Biol, 24(1), 87–97. https://doi.org/10.1016/j.chembiol.2016.12.007

Jiang, H., Zhang, X., Chen, X., Aramsangtienchai, P., Tong, Z., & Lin, H. (2018). Protein Lipidation: Occurrence, Mechanisms, Biological Functions, and Enabling Technologies. Chem Rev, 118(3), 919–988. https://doi.org/10.1021/acs.chemrev.6b00750

Karmi, A., Iozzo, P., Viljanen, A., Hirvonen, J., Fielding, B. A., Virtanen, K., Oikonen, V., Kemppainen, J., Viljanen, T., Guiducci, L., Haaparanta-Solin, M., Nagren, K., Solin, O., & Nuutila, P. (2010). Increased brain fatty acid uptake in metabolic syndrome. Diabetes, 59(9), 2171–2177. https://doi.org/10.2337/db09-0138

Kathayat, R. S., Elvira, P. D., & Dickinson, B. C. (2017). A fluorescent probe for cysteine depalmitoylation reveals dynamic APT signaling. Nat Chem Biol, 13(2), 150–152. https://doi.org/10.1038/nchembio.2262

Katz, M., Amit, I., & Yarden, Y. (2007). Regulation of MAPKs by growth factors and receptor tyrosine kinases. Biochim Biophys Acta, 1773(8), 1161–1176. https://doi.org/10.1016/j.bbamcr.2007.01.002

Ke, M., Yuan, X., He, A., Yu, P., Chen, W., Shi, Y., Hunter, T., Zou, P., & Tian, R. (2021). Spatiotemporal profiling of cytosolic signaling complexes in living cells by selective proximity proteomics. Nat Commun, 12(1), 71. https://doi.org/10.1038/s41467-020-20367-x

Kiyatkin, A., van Alderwerelt van Rosenburgh, I. K., Klein, D. E., & Lemmon, M. A. (2020). Kinetics of receptor tyrosine kinase activation define ERK signaling dynamics. Sci Signal, 13(645). https://doi.org/10.1126/scisignal.aaz5267

Kutzleb, C., Sanders, G., Yamamoto, R., Wang, X., Lichte, B., Petrasch-Parwez, E., & Kilimann, M. W. (1998). Paralemmin, a prenyl-palmitoyl-anchored phosphoprotein abundant in neurons and implicated in plasma membrane dynamics and cell process formation. J Cell Biol, 143(3), 795–813. https://doi.org/10.1083/jcb.143.3.795

Lake, D., Correa, S. A., & Muller, J. (2016). Negative feedback regulation of the ERK1/2 MAPK pathway. Cell Mol Life Sci, 73(23), 4397–4413. https://doi.org/10.1007/s00018-016-2297-8

Lan, T., Delalande, C., & Dickinson, B. C. (2021). Inhibitors of DHHC family proteins. Curr Opin Chem Biol, 65, 118–125. https://doi.org/10.1016/j.cbpa.2021.07.002

Lavoie, H., Gagnon, J., & Therrien, M. (2020). ERK signalling: a master regulator of cell behaviour, life and fate. Nat Rev Mol Cell Biol, 21(10), 607–632. https://doi.org/10.1038/s41580-020-0255-7

Lin, D. T., & Conibear, E. (2015). ABHD17 proteins are novel protein depalmitoylases that regulate N-Ras palmitate turnover and subcellular localization. Elife, 4, e11306. https://doi.org/10.7554/eLife.11306

Lu, N., & Malemud, C. J. (2019). Extracellular Signal-Regulated Kinase: A Regulator of Cell Growth, Inflammation, Chondrocyte and Bone Cell Receptor-Mediated Gene Expression. Int J Mol Sci, 20(15). https://doi.org/10.3390/ijms20153792

Miao, L., & Tian, H. (2020). Development of ERK1/2 inhibitors as a therapeutic strategy for tumour with MAPK upstream target mutations. J Drug Target, 28(2), 154–165. https://doi.org/10.1080/1061186X.2019.1648477

Morrison, D. K. (2012). MAP kinase pathways. Cold Spring Harb Perspect Biol, 4(11). https://doi.org/10.1101/cshperspect.a011254

Niphakis, M. J., Lum, K. M., Cognetta, A. B., 3rd, Correia, B. E., Ichu, T. A., Olucha, J., Brown, S. J., Kundu, S., Piscitelli, F., Rosen, H., & Cravatt, B. F. (2015). A Global Map of Lipid-Binding Proteins and Their Ligandability in Cells. Cell, 161(7), 1668–1680. https://doi.org/10.1016/j.cell.2015.05.045

Ohno, Y., Kihara, A., Sano, T., & Igarashi, Y. (2006). Intracellular localization and tissue-specific distribution of human and yeast DHHC cysteine-rich domain-containing proteins. Biochim Biophys Acta, 1761(4), 474–483. https://doi.org/10.1016/j.bbalip.2006.03.010

Oppermann, F. S., Gnad, F., Olsen, J. V., Hornberger, R., Greff, Z., Keri, G., Mann, M., & Daub, H. (2009). Large-scale proteomics analysis of the human kinome. Mol Cell Proteomics, 8(7), 1751–1764. https://doi.org/10.1074/mcp.M800588-MCP200

Ozaki, K. I., Awazu, M., Tamiya, M., Iwasaki, Y., Harada, A., Kugisaki, S., Tanimura, S., & Kohno, M. (2016). Targeting the ERK signaling pathway as a potential treatment for insulin resistance and type 2 diabetes. Am J Physiol Endocrinol Metab, 310(8), E643–E651. https://doi.org/10.1152/ajpendo.00445.2015

Papa, S., Choy, P. M., & Bubici, C. (2019). The ERK and JNK pathways in the regulation of metabolic reprogramming. Oncogene, 38(13), 2223–2240. https://doi.org/10.1038/s41388-018-0582-8

Peti, W., & Page, R. (2013). Molecular basis of MAP kinase regulation. Protein Sci, 22(12), 1698–1710. https://doi.org/10.1002/pro.2374

Pinilla-Macua, I., Grassart, A., Duvvuri, U., Watkins, S. C., & Sorkin, A. (2017). EGF receptor signaling, phosphorylation, ubiquitylation and endocytosis in tumors in vivo. Elife, 6. https://doi.org/10.7554/eLife.31993

Plotnikov, A., Chuderland, D., Karamansha, Y., Livnah, O., & Seger, R. (2019). Nuclear ERK Translocation is Mediated by Protein Kinase CK2 and Accelerated by Autophosphorylation. Cell Physiol Biochem, 53(2), 366–387. https://doi.org/10.33594/000000144

Plotnikov, A., Flores, K., Maik-Rachline, G., Zehorai, E., Kapri-Pardes, E., Berti, D. A., Hanoch, T., Besser, M. J., & Seger, R. (2015). The nuclear translocation of ERK1/2 as an anticancer target. Nat Commun, 6, 6685. https://doi.org/10.1038/ncomms7685

Qiu, T., Kathayat, R. S., Cao, Y., Beck, M. W., & Dickinson, B. C. (2018). A Fluorescent Probe with Improved Water Solubility Permits the Analysis of Protein S-Depalmitoylation Activity in Live Cells. Biochemistry, 57(2), 221–225. https://doi.org/10.1021/acs.biochem.7b00835

Raman, M., Chen, W., & Cobb, M. H. (2007). Differential regulation and properties of MAPKs. Oncogene, 26(22), 3100–3112. https://doi.org/10.1038/sj.onc.1210392

Ren, W., Jhala, U. S., & Du, K. (2013). Proteomic analysis of protein palmitoylation in adipocytes. Adipocyte, 2(1), 17–28. https://doi.org/10.4161/adip.22117

Roskoski, R., Jr. (2012). ERK1/2 MAP kinases: structure, function, and regulation. Pharmacol Res, 66(2), 105–143. https://doi.org/10.1016/j.phrs.2012.04.005

Runkle, K. B., Kharbanda, A., Stypulkowski, E., Cao, X. J., Wang, W., Garcia, B. A., & Witze, E. S. (2016). Inhibition of DHHC20-Mediated EGFR Palmitoylation Creates a Dependence on EGFR Signaling. Mol Cell, 62(3), 385–396. https://doi.org/10.1016/j.molcel.2016.04.003

Sadeghi, R. S., Kulej, K., Kathayat, R. S., Garcia, B. A., Dickinson, B. C., Brady, D. C., & Witze, E. S. (2018). Wnt5a signaling induced phosphorylation increases APT1 activity and promotes melanoma metastatic behavior. Elife, 7. https://doi.org/10.7554/eLife.34362

Sebolt-Leopold, J. S., Dudley, D. T., Herrera, R., Van Becelaere, K., Wiland, A., Gowan, R. C., Tecle, H., Barrett, S. D., Bridges, A., Przybranowski, S., Leopold, W. R., & Saltiel, A. R. (1999). Blockade of the MAP kinase pathway suppresses growth of colon tumors in vivo. Nat Med, 5(7), 810–816. https://doi.org/10.1038/10533

Shaul, Y. D., & Seger, R. (2007). The MEK/ERK cascade: from signaling specificity to diverse functions. Biochim Biophys Acta, 1773(8), 1213–1226. https://doi.org/10.1016/j.bbamcr.2006.10.005

Spinelli, M., Fusco, S., & Grassi, C. (2018). Nutrient-Dependent Changes of Protein Palmitoylation: Impact on Nuclear Enzymes and Regulation of Gene Expression. Int J Mol Sci, 19(12). https://doi.org/10.3390/ijms19123820

Sun, J., & Nan, G. (2017). The extracellular signal-regulated kinase 1/2 pathway in neurological diseases: A potential therapeutic target (Review). Int J Mol Med, 39(6), 1338–1346. https://doi.org/10.3892/ijmm.2017.2962

Sun, Y., Liu, W. Z., Liu, T., Feng, X., Yang, N., & Zhou, H. F. (2015). Signaling pathway of MAPK/ERK in cell proliferation, differentiation, migration, senescence and apoptosis. J Recept Signal Transduct Res, 35(6), 600–604. https://doi.org/10.3109/10799893.2015.1030412

Tai, W. M., Yong, W. P., Lim, C., Low, L. S., Tham, C. K., Koh, T. S., Ng, Q. S., Wang, W. W., Wang, L. Z., Hartano, S., Thng, C. H., Huynh, H., Lim, K. T., Toh, H. C., Goh, B. C., & Choo, S. P. (2016). A phase Ib study of selumetinib (AZD6244, ARRY-142886) in combination with sorafenib in advanced hepatocellular carcinoma (HCC). Ann Oncol, 27(12), 2210–2215. https://doi.org/10.1093/annonc/mdw415

Tsai, J., Lee, J. T., Wang, W., Zhang, J., Cho, H., Mamo, S., Bremer, R., Gillette, S., Kong, J., Haass, N. K., Sproesser, K., Li, L., Smalley, K. S., Fong, D., Zhu, Y. L., Marimuthu, A., Nguyen, H., Lam, B., Liu, J., Cheung, I., Rice, J., Suzuki, Y., Luu, C., Settachatgul, C., Shellooe, R., Cantwell, J., Kim, S. H., Schlessinger, J., Zhang, K. Y., West, B. L., Powell, B., Habets, G., Zhang, C., Ibrahim, P. N., Hirth, P., Artis, D. R., Herlyn, M., & Bollag, G. (2008). Discovery of a selective inhibitor of oncogenic B-Raf kinase with potent antimelanoma activity. Proc Natl Acad Sci U S A, 105(8), 3041–3046. https://doi.org/10.1073/pnas.0711741105

Uhlitz, F., Sieber, A., Wyler, E., Fritsche-Guenther, R., Meisig, J., Landthaler, M., Klinger, B., & Bluthgen, N. (2017). An immediate-late gene expression module decodes ERK signal duration. Mol Syst Biol, 13(5), 928. https://doi.org/10.15252/msb.20177554

Vougiouklakis, T., Sone, K., Saloura, V., Cho, H. S., Suzuki, T., Dohmae, N., Alachkar, H., Nakamura, Y., & Hamamoto, R. (2015). SUV420H1 enhances the phosphorylation and transcription of ERK1 in cancer cells. Oncotarget, 6(41), 43162–43171. https://doi.org/10.18632/oncotarget.6351

Wang, C. Y., & Liao, J. K. (2012). A mouse model of diet-induced obesity and insulin resistance. Methods Mol Biol, 821, 421–433. https://doi.org/10.1007/978-1-61779-430-8_27

Webb, Y., Hermida-Matsumoto, L., & Resh, M. D. (2000). Inhibition of protein palmitoylation, raft localization, and T cell signaling by 2-bromopalmitate and polyunsaturated fatty acids. J Biol Chem, 275(1), 261–270. https://doi.org/10.1074/jbc.275.1.261

Wortzel, I., & Seger, R. (2011). The ERK Cascade: Distinct Functions within Various Subcellular Organelles. Genes Cancer, 2(3), 195–209. https://doi.org/10.1177/1947601911407328

Wu, J. Y., Xiang, S., Zhang, M., Fang, B., Huang, H., Kwon, O. K., Zhao, Y., Yang, Z., Bai, W., Bepler, G., & Zhang, X. M. (2018). Histone deacetylase 6 (HDAC6) deacetylates extracellular signal-regulated kinase 1 (ERK1) and thereby stimulates ERK1 activity. J Biol Chem, 293(6), 1976–1993. https://doi.org/10.1074/jbc.M117.795955

Yang, G., Liu, Y., Yang, K., Liu, R., Zhu, S., Coquinco, A., Wen, W., Kojic, L., Jia, W., & Cynader, M. (2012). Isoform-specific palmitoylation of JNK regulates axonal development. Cell Death Differ, 19(4), 553–561. https://doi.org/10.1038/cdd.2011.124

Zhang, M., Zhou, L., Xu, Y., Yang, M., Xu, Y., Komaniecki, G. P., Kosciuk, T., Chen, X., Lu, X., Zou, X., Linder, M. E., & Lin, H. (2020). A STAT3 palmitoylation cycle promotes TH17 differentiation and colitis. Nature, 586(7829), 434–439. https://doi.org/10.1038/s41586-020-2799-2

